# Prefrontal projections modulate recurrent circuitry in insular cortex to support short-term memory

**DOI:** 10.1101/2023.04.28.538791

**Authors:** Jian Yao, Ruiqing Hou, Hongmei Fan, Jincan Hou, Qi Cheng, Chengyu T. Li

## Abstract

Short-term memory (STM) maintains information during a short delay period. How long-range and local connections interact to support STM encoding remains elusive. Here we tackled the problem focusing on long-range projections from medial prefrontal cortex (mPFC) to anterior agranular insular cortex (aAIC) in head-fixed mice performing an olfactory delayed-response task. Optogenetic and electrophysiological experiments revealed behavioral importance of the two regions in encoding STM information. Spike-correlogram analysis revealed strong local and cross-region functional coupling (FC) between memory neurons encoding the same information. Optogenetic suppression of mPFC-aAIC projections during the delay period reduced behavioral performance, the proportion of memory neurons and memory-specific FC within aAIC, whereas optogenetic excitation enhanced all of them. MPFC-to-aAIC projections also bidirectionally modulated efficacy of STM-information transfer, measured by contribution of FC spiking pairs to coding ability of following neurons. Thus, prefrontal projections modulate functional connectivity and coding ability of insular neurons to support short-term memory.

## INTRODUCTION

Short-term memory (STM)^1, 2^ actively maintains information for a short delay period, which is critical for linking the immediate past with current demands to achieve appropriate behavioral outcome. Internally generated neuronal activity capable of encoding the maintained STM information is necessary for STM behavior and has been widely observed in various regions^3–8^. Causal perturbations using lesion, pharmacological or optogenetic approaches have demonstrated the functional importance of the neurons in many regions for STM maintenance, including sensory, prefrontal and premotor regions^5, 9–13^. Recently, pathway-specific perturbation has demonstrated the importance of long-range connection for STM, such as entorhinal cortex to hippocampus, hippocampus-to-mPFC and mediodorsal thalamus to mPFC projections for spatial STM^14–16^, thalamus to anterior lateral motor cortex (ALM) and ALM to cerebellum for motor-oriented STM^10, 17^, and mPFC to aAIC for sensory-oriented STM tasks^18^. However, neural mechanisms underlying STM modulation by cross-region projections remain unclear.

Here we focused on what kind of local-circuit processing supporting STM is modulated by long-range connections. Brain functions rely on recurrent connectivity, which could link neurons encoding the same information into cell assembly^19–22^ through the fire-together-wire-together principle^19^. Correlation in neuronal spiking reveals dynamic patterns of recurrent connectivity^23^ and is related with synaptic connection^24^. In particular, correlated spiking at millisecond temporal scale could reveal directional functional coupling (FC)^25–28^, which was associated with various behaviors^22, 29–34^. For example, the neurons in lateral prefrontal cortex (LPFC) have been shown to exhibit stronger correlation among the neurons similarly tuned to STM information^35, 36^. However, it remains unknown how STM-related FC is controlled by long-range cortical projections.

To address the question, we trained mice to perform a delayed-response STM task and applied optogenetics and electrophysiological recording to mPFC and aAIC, two regions critical for the STM task performance and information maintenance during the learning phase. Importantly, pathway-specific optogenetic manipulation of the delay-period activity of mPFC projections to aAIC induced bidirectional modulation in behavioral performance, as well as the coding ability and local FC of aAIC memory neurons. In-depth analysis showed that the efficacy of information transfer, defined by the contribution of functional coupled spiking events to the coding ability of the following neurons in a FC neuronal pair, was also bidirectionally modulated by mPFC-aAIC projections. Thus, long-range projections from prefrontal cortex are critical in controlling memory coding ability and functional coupling of insular neurons to support STM behavior.

## RESULTS

### Importance of Delay-Period Activity of Both mPFC and aAIC Neurons for STM

We trained head-fixed mice to perform an olfactory delayed-response task (DRT) using an automatic training system^37^. In each trial of the motor-oriented DRT (Figures 1A3 and 1F3), mice made Go or NoGo decision based on the identity of odor sample (propyl formate for Go, 1-butanol for NoGo decision; sample-odor delivery duration: 1 sec). Mice needed to withhold licking during a delay period of 4 or 10 sec, otherwise the trials would be abolished without reward (Figures S1E and S1H). The DRT delay-period activity could reflect sensory memory, reward prediction, motor planning^38, 39^ or timing expectation^40–42^. Mice readily learnt the task, as reflected by the increased correct rejection (CR) rate, discriminability (*d’*), and lick efficiency throughout the training (Figures 1B, 1G, S1C, S1D, S1F, and S1G). There was no change in the hit rate throughout learning (Figures 1B and 1G).

**Figure 1.**
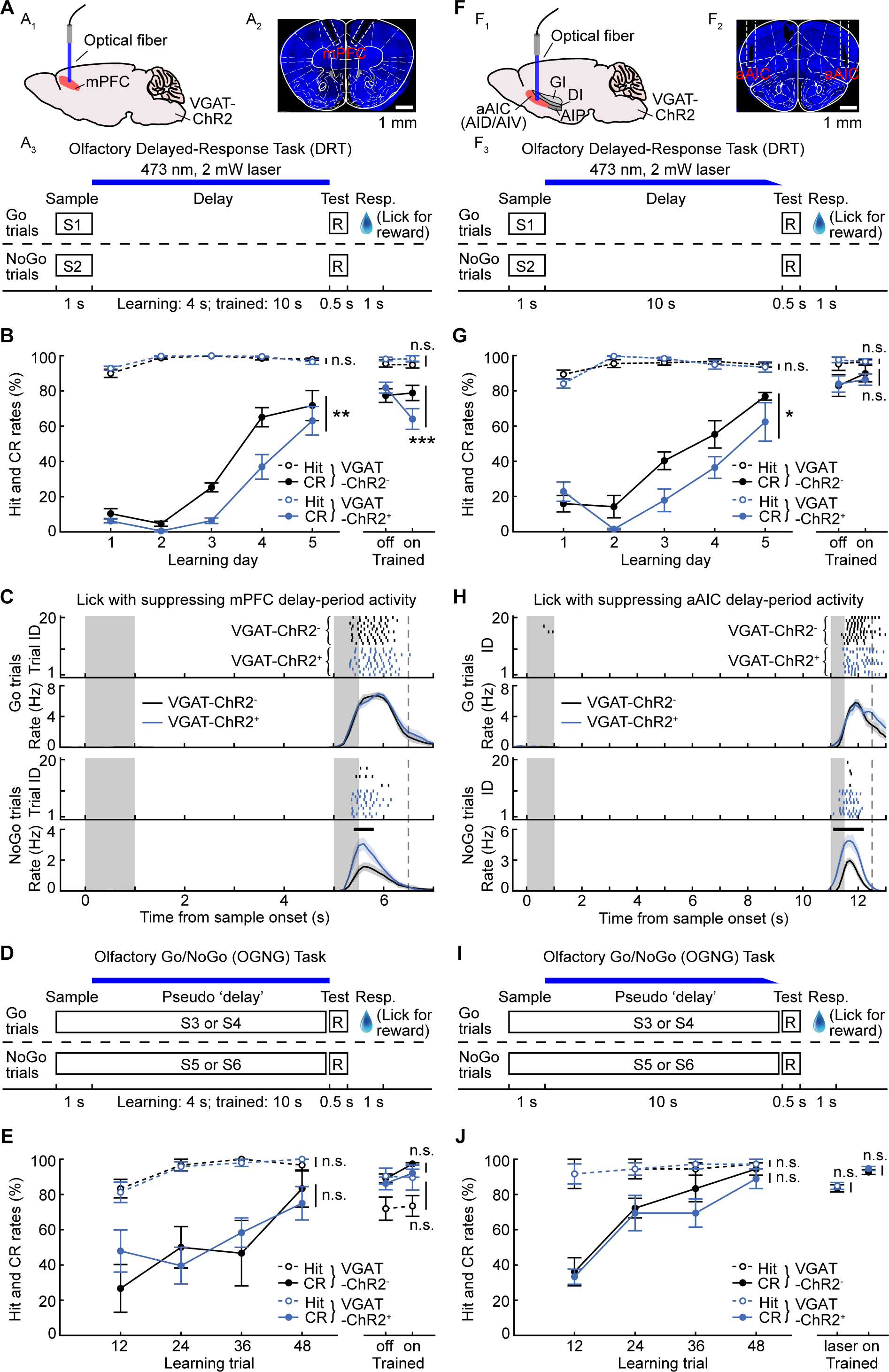
Importance of delay-period activity of both mPFC and aAIC neurons for STM. (A) Schematic diagram (A1), histology image of optical fibers (A2), and behavioral design (A3) for the mPFC optogenetic suppression experiment in the delayed-response task (DRT). S1, propyl formate; S2, 1-butanol; R, heptane. (B) Hit (broken lines) and correct-rejection (CR, solid lines) rates of mice in performing DRT, for optogenetic suppression of the mPFC delay-period activity (blue) and control (black) groups. For the hit rate, learning day: main effect of genotypes F (1,19) = 0.87, p = 0.36; trained: p = 0.49 for laser off/on. For the CR rate, learning day: F (1,19) = 12.80, p = 0.002; trained: p = 6.22 × 10^−4^ for laser off/on. Statistics: Two-way ANOVA with mixed design (Tw-ANOVA-md) for learning-day performance, n = 11 and 10 mice for VGAT-ChR2^+^ and VGAT-ChR2^−^ groups, respectively; Mann-Whitney *U* test for the well-trained phase, n = 8 mice for both groups. Error bars represent mean ± SEM. (C) Lick raster and lick rate for all mice in Go and NoGo trials. Note the absence of lick during the delay period. Statistics: p < 0.05, cluster-based permutation test. Shaded area represents mean ± SEM. (D) Diagram for the task design of the OGNG task. Note the presence of sample odor during the pseudo ‘delay’ period. S3, butyl formate; S4, ethyl acetate; S5, 1-pentanol; S6, 3-methyl-2-buten-1-ol; R, heptane. (E) Hit and CR rates of mice in performing OGNG task, for optogenetic suppression of mPFC activity during the ‘pseudo’ delay period (blue) and control (black) groups. For the hit rate, learning day: F (1,11) = 0.022, p = 0.88; trained: p = 0.39 for laser off/on. For the CR rate, learning day: F (1,11) = 0.074, p = 0.79; trained: p = 0.33 for laser off/on. Statistics: Tw-ANOVA-md for learning-day performance, n = 8 and 5 mice for VGAT-ChR2+ and VGAT-ChR2-groups, respectively; Mann-Whitney U test for the well-trained phase, n = 6 mice for both groups. (F) As (A) for suppressing aAIC delay-period activity. Optical fibers were implanted into aAIC (AID/AIV in the Allen Brain Atlas) bilaterally. aAIC, anterior agranular insular cortex; GI, granular insular cortex; DI, dysgranular insular cortex; AIP, posterior agranular insular cortex. (G) As (B) for suppression of aAIC. For the hit rate, learning day: F (1,13) = 0.042, p = 0.84; trained: p = 0.79 for laser off/on. For the CR rate, learning day: F (1,13) = 6.30, p = 0.026; trained: p = 0.66 for laser off/on. Statistics: Tw-ANOVA-md for learning-day performance, n = 7 and 8 mice for VGAT-ChR2^+^ and VGAT-ChR2^−^ groups, respectively; Mann-Whitney *U* test for the well-trained phase, n = 8 mice for both groups. (H) As (C) for suppressing aAIC delay-period activity. (I) As (D) for suppression of aAIC. (J) As (E) for suppressing aAIC activity. For the hit rate, learning day: F (1,10) = 0.017, p = 0.90; trained: p = 1.00. For the CR rate, learning day: F (1,10) = 2.58, p = 0.14; trained: p = 0.80. Statistics: Tw-ANOVA-md for learning-day performance, Mann-Whitney U test for the well-trained phase. n = 6 mice for both groups.

See also Figure S1.

We then examined the role of the delay-period activity of mPFC and aAIC in performing this STM task. The mPFC is a crucial region for STM, as shown by behavioral impairment following perturbation of PFC activities in non-human primates and rodents^9, 43–46^. The delay-period activity of aAIC has also been shown to be important for successful maintenance of sensory-oriented STM in the learning phase^18^. To causally examine the importance of delay-period activity of either mPFC or aAIC in learning or well-trained phase of motor-oriented DRT, we activated local inhibitory neurons in these regions to suppress the activity of the excitatory neurons. Transgenic mice expressing channel-rhodopsin (ChR2) only in the GABAergic neurons [with the promoter of vesicular GABA transporter (VGAT), ref.^47^; referred to as VGAT-ChR2^+^] were used for optogenetic manipulation, with the ChR2-negative litter mates (VGAT-ChR2^−^) as control in a blind design^6, 18^. Optical fiber was bilaterally inserted on top of mPFC (Figures 1A1, 1A2, and S1J) or aAIC (Figures 1F1, 1F2, and S1K) and the efficiency of optogenetic suppression was verified with extracellular op-tetrodes recording (Figures S1A and S1B). This method has been extensively used and proven to be efficient for deciphering the causal contribution of excitatory neurons in various behavioral designs^48–50^.

Optogenetic suppression of delay-period activities in either mPFC or aAIC significantly decreased the CR rate, discriminability (*d’*), and lick efficiency during the first five days of learning to perform the DRT (Figures 1B, 1C, 1G, 1H, S1C, S1D, S1F, and S1G), without changing the hit rate and the proportion of aborted trials (Figures 1B, 1C, 1G, 1H, S1E, and S1H). We then suppressed the delay-period activities of mPFC or aAIC in the well-trained phase (averaged performance correct rate for consecutive 60 trials > 80%). Optogenetic suppression of delay-period activities in mPFC but not aAIC significantly decreased the CR rate, discriminability (*d’*), and lick efficiency during the well-trained phase (Figure S1I and right panels of Figures 1B, 1G, S1C, S1D, S1F, and S1G).

Besides STM, other diverse brain functions could involve mPFC^51^ and aAIC^52^, including sensory processing^53–58^, attention^59–61^, motivation^62, 63^, decision making^64^, reward expectation^65, 66^, motor control^67, 68^, reward valence^69, 70^, and learning^71, 72^. To further control for such non-STM processes, we designed a control experiment of olfactory Go/NoGo (OGNG) task. Two sample odors were designed as Go cues (butyl formate and ethyl acetate) and two other odors as NoGo cues (1-pentanol and 3-methyl-2-buten-1-ol). The OGNG task structure is similar to that of DRT, with OGNG sample odors delivered during the corresponding delay period of DRT (Figures 1D and 1I). Thus, there is no need to maintain STM in the OGNG task, but the abovementioned non-STM processes should still be involved. If suppression of delay-period activity in mPFC or aAIC doesn’t produce the behavioral deficit, then it is less likely that impaired DRT performance was due to the abovementioned non-STM processes. As shown in Figures 1E and 1J, suppressing the neuronal activities of the pseudo ‘delay’ period in neither mPFC nor aAIC impaired performance of the OGNG task. In the DRT, lick rates in the Go trials (∼100% hit rate for both groups, see Figures 1B and 1G) and the aborted trials proportion (Figures S1E and S1H) remained unchanged following optogenetic suppression of mPFC or aAIC, therefore the impaired performance was not due to motor action. Thus, optogenetics-induced impairment in DRT performance (Figures 1B and 1G) is not likely to be due to the non-STM processes mentioned above.

In summary, the delay-period activities of both mPFC and aAIC neurons are critical for maintaining STM information in the motor-oriented DRT, especially in the learning phase.

### Encoding STM information by aAIC and mPFC Neurons

Spiking activity of both persistent^12, 17, 73–80^ and transient^18, 81–97^ patterns during the delay period could be the neural underpinning of STM. On the one hand, recurrent network dynamics by point attractor could support persistent activity, for example in ALM^12, 17^ and mPFC^44, 45, 78^ neurons. On the other hand, population activity waves relayed through transiently active neurons could also encode STM information^87–89^, the functional relevance of which to olfactory STM was demonstrated by simultaneous optogenetic perturbation and electrophysiological recordings in aAIC neurons^18^. We hereby examined the neuronal dynamics in mPFC and aAIC in the motor-oriented STM task.

To examine the dynamic patterns of DRT associated-neural activity in the learning phase, we recorded single-unit activity in both aAIC and mPFC using custom-made tetrodes (electrode as in ref.^6, 18^; schematic diagram of aAIC and mPFC recordings in Figures 2A and 2K, respectively). The recording sites for the example neurons are shown in Figures 2B and 2L and all neurons in S2A (quality control of single-unit recording in Figures S2B-S2G). Recordings began on the first day of DRT training, with daily advancing the recording electrodes (about 50 μm/day). We obtained 816 mPFC neurons and 437 aAIC neurons (Figure 2) from 8 mice in the first 6 days of learning. Activity of many neurons exhibited delay-period modulation of diverse patterns, as shown by the raster plot of example neurons (Figures 2C, 2D, 2M, and 2N) and activity heatmap of all neurons (Figures 2E and 2O).

**Figure 2.**
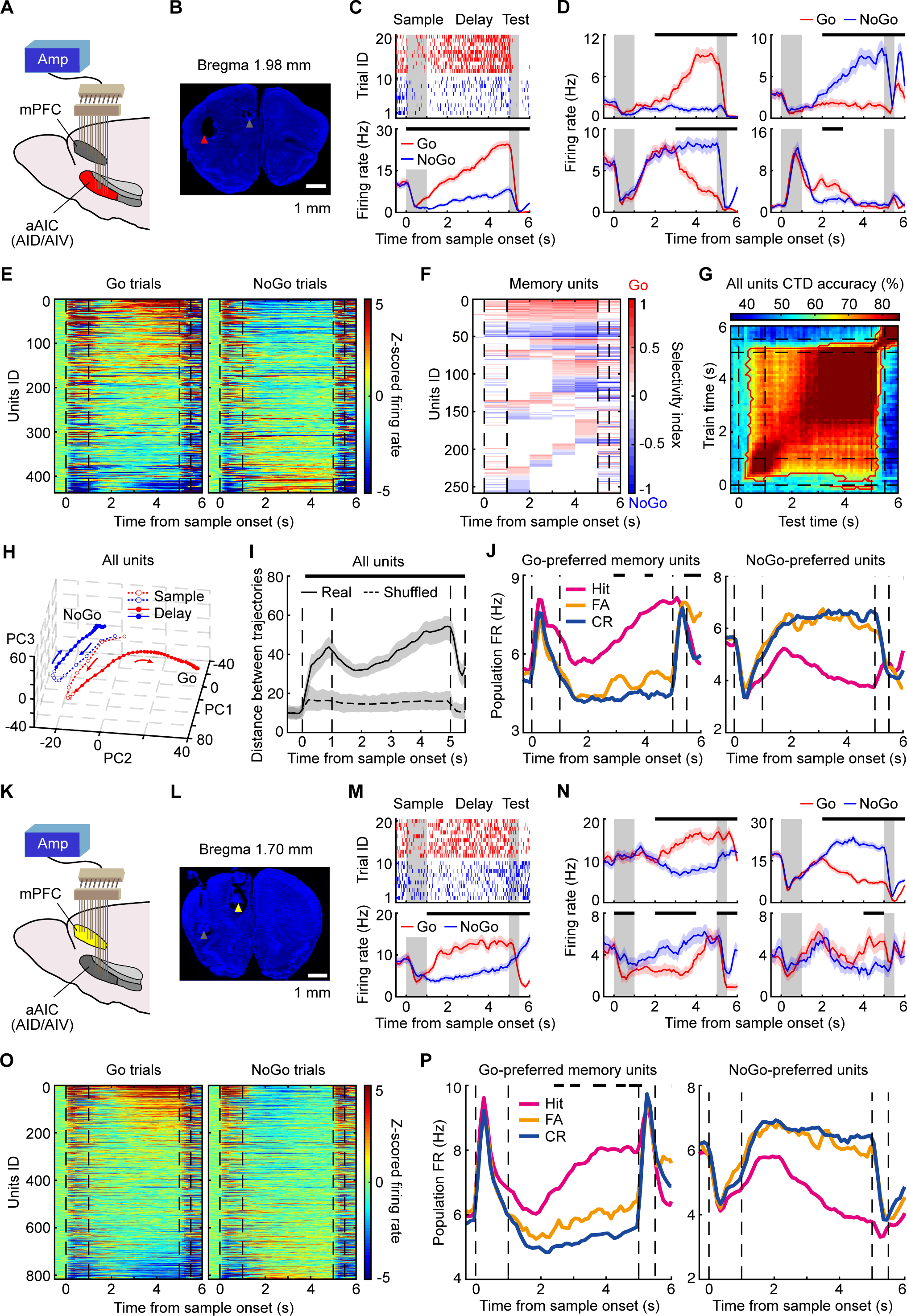
Encoding of STM information by aAIC and mPFC neurons. (A) Diagram for single-unit recording of aAIC activity. (B) Histology for unilateral-recording sites in aAIC. (C) Raster plot and peristimulus time histogram (PSTH) showing the activity of an example aAIC neuron in DRT. Separate black bars on top indicated the time bins (size: 1 sec) with statistically significant selectivity for two curves throughout all figures. Statistics: p < 0.05, Mann-Whitney *U* test with Bonferroni correction. Shaded area represents mean ± SEM. (D) PSTH for other four example aAIC neurons. (E) Heatmaps of aAIC population activity following different sample odors. Each row of the two heatmaps was from the same neuron. (F) Heatmap plot showing the memory-selectivity profiles of aAIC memory neurons. (G) Cross-temporal decoding (CTD) results with all neurons in aAIC. Red contour outlined the time points with significant decoding accuracy. Statistics: p < 0.05, cluster-based permutation test. (H) aAIC population activity trajectories based on the first three principal components (explaining 83% variance in PCA), following different sample odors (bin size: 100 msec). Arrow indicated the time course. (I) Distance in PCA trajectories based on the first 20 PC. Shadow indicates 95% confidence interval (CI) from bootstrapping (100 and 1000 runs for the real and shuffled data, respectively). Statistics: p < 0.05, cluster-based permutation test. (J) Averaged activities of Go-preferred (left, n = 35 neurons) and NoGo-preferred (right, n = 38 neurons) aAIC neurons in hit, false alarm (FA), and CR trials. Statistics: p < 0.05, cluster-based permutation test. (K) Diagram for single-unit recording of mPFC activity. (L) Histology for unilateral-recording sites in mPFC. (M) As (C) for an example mPFC neuron. (N) As (D) for other four example mPFC neurons. (O) As (E) for mPFC. (P) As (J) for Go-preferred (left, n = 54 neurons) and NoGo-preferred (right, n = 51 neurons) mPFC neurons.

See also Figures S2-S4.

The delay-period activity of many mPFC and aAIC neurons carried the maintained STM information, which was reflected by the differential activities following different sample odors (examples in Figures 2C, 2D, 2M, and 2N; activity heatmap of all recorded mPFC and aAIC neurons in Figures 2E and 2O). These neurons were defined as the ‘memory neurons’ and could be classified into three groups (as in ref.^81, 87^, see **Methods**): (1) transient-encoding memory neurons showing selectivity for 1-3 sec (p < 0.05, Bonferroni multiple comparison corrected Mann-Whitney *U* test, unless stated otherwise; see Figures 2D, 2F, and 2N), (2) sustained-encoding memory neurons showing selectivity within all 4 one-second delay bins (Figures 2C, 2F, and 2M), (3) un-classified memory neurons (37 out of 816 for mPFC, 18 out of 437 for aAIC, see Figures S3A and S3B). Other neurons were not selective to STM information in any time bin of the delay period (54.29% in mPFC and 40.96% in aAIC). The sustained neurons were fewer than the transient neurons (7.84% *vs.* 37.87% in Figure S3C for mPFC, 11.44% vs. 47.60% in Figure S3F for aAIC, p < 0.001, chi-square test). The timing of significant selectivity of transient neurons tiled over the entire delay period (Figure 2F). The potential bias of statistical definition in transient and sustained neurons was excluded as in the previous study^18^ (see **Methods**).

At single-trial level, the spiking activities of most, if not all, sustained neurons were not as ‘persistent’ as averaged peristimulus time histogram (PSTH, reflected by the overlapped activities between different sample odors, see Figures S3I-S3K), suggesting that the cross-trial averaging might contribute to the observed sustained firing patterns in our current recordings (Figure S3L, as in ref.^18^).

To further test the stability of STM-encoding ability of mPFC and aAIC neurons in information maintenance during the DRT delay period, we performed the population cross-temporal decoding (CTD) analysis^85, 86^. The essence of this analysis is to use the activity at a given time to decode the maintained information at other time points, with systematic changes in the timing of both training and testing data. A significant decoding power in the off-diagonal space of a CTD plot suggests more persistent pattern for information maintenance. Indeed, we observed significant off-diagonal decoding power for the recorded aAIC neurons (Figure 2G, CTD result for all mPFC neurons not shown). Furthermore, both sustained and transient neurons exhibited stable CTD ability, with higher persistence in encoding STM information for sustained neurons (mPFC in Figures S3D and S3E, aAIC in Figures S3G and S3H).

To further examine and visualize patterns of population dynamics, we performed dimensionality reduction analysis by principal component analysis (PCA) and revealed clear separation in population trajectories of aAIC population activities evoked by Go vs. NoGo odors (trajectories in Figure 2H; distance measures in Figure 2I; PCA result for mPFC not shown). Thus, downstream neurons could stably readout STM information from population activities of mPFC and aAIC neurons, which constituted separated trajectories on neural manifold encoding STM information.

We further examined the relationship between neuronal firing and behavioral performance. During the sample-delivery period, there was no difference among the neuronal activities in the hit, CR, and false alarm (FA) trials (aAIC and mPFC in Figures 2J and 2P, respectively). For the Go-preferred neurons, the activity of the FA trials fell between that of the hit and CR trials, leaning toward the CR trials (left panels of Figures 2J and 2P for aAIC and mPFC, respectively). Noted the prominent ramping-up activity of delay period in the hit trials for the Go-preferred neurons. Such ramping activity was previously observed in many premotor and prefrontal regions^12, 17, 79, 80^ and believed to be important for motor preparation^38, 39^ or prediction of timing^40–42^.

For the NoGo-preferred neurons, there was no difference between the activities of the FA and CR trials (right panels of Figures 2J and 2P for aAIC and mPFC, respectively). Contrary to Go-preferred neurons, delay-period activity of NoGo-preferred neurons exhibited evident ramping-down patterns in the hit trials. The obvious ramping activity of delay period in FA or CR trials was not observed for both Go-preferred and NoGo-preferred neurons (Figures 2J and 2P). Therefore, the delay-period activities of both mPFC and aAIC were correlated with behavioral performance, but only for the Go-preferred neurons.

After the delay period, the activities of memory neurons in the hit, CR, and FA trials converged to similar levels following the application of response cue (Figures 2J and 2P). This phenomenon suggested that the memory neurons were not responsible for Go/NoGo action *per se*. We indeed observed that most memory neurons lost or switched their memory-coding ability during the action period (example neurons in Figures S4A-S4D, proportion in Figures S4E and S4F). Consistently, we observed sharp decline then increase in off-diagonal decoding following the application of response cue in the CTD plots (Figures 2G, S3D, and S3G), suggesting that different subpopulations of neurons were differentially involved in the delay and action periods.

Taken together, our results showed that the aAIC and mPFC neurons employ both transient and sustained patterns during the delay period to encode the motor-oriented STM-information, in a manner correlated with behavioral performance.

### Preferential Functional Coupling between Memory Neurons

To test whether the neurons encoding the same information are strongly connected as predicted by the cell-assembly hypothesis^19–22^, we measured functional coupling (FC) using spike-correlogram analysis^98–101^ between simultaneously recorded neurons. The shift-predictor-subtracted cross-correlogram (SSCC)^98–101^ of two example neuronal pairs (one pair within mPFC in Figure 3A, the other within aAIC in Figure 3B) showed a prominent peak on the right side of SSCC (p < 0.05, within 10 msec after spiking of a leading neuron, Student *t*-test with Bonferroni correction compared with trials-shuffled spiking data; more examples in Figures S5A-S5F). The short latency (<10 msec) and asymmetry of the correlogram profile indicated directional FC from a leading neuron to a following neuron (Figures 3A3 and 3B3). To control for SSCC peaks due to common inputs, we excluded the neuronal pairs with significantly correlated spiking within -2 to +2 msec of leading-neuron spikes, as in ref.^102^. In FC neuronal pairs, the functionally coupled spiking pairs (FCSPs) events were defined as correlated firing of both leading and following neurons with 2-10 msec intervals (thick line pairs in Figures 3A2 and 3B2). We then searched FC within and across mPFC and aAIC.

**Figure 3.**
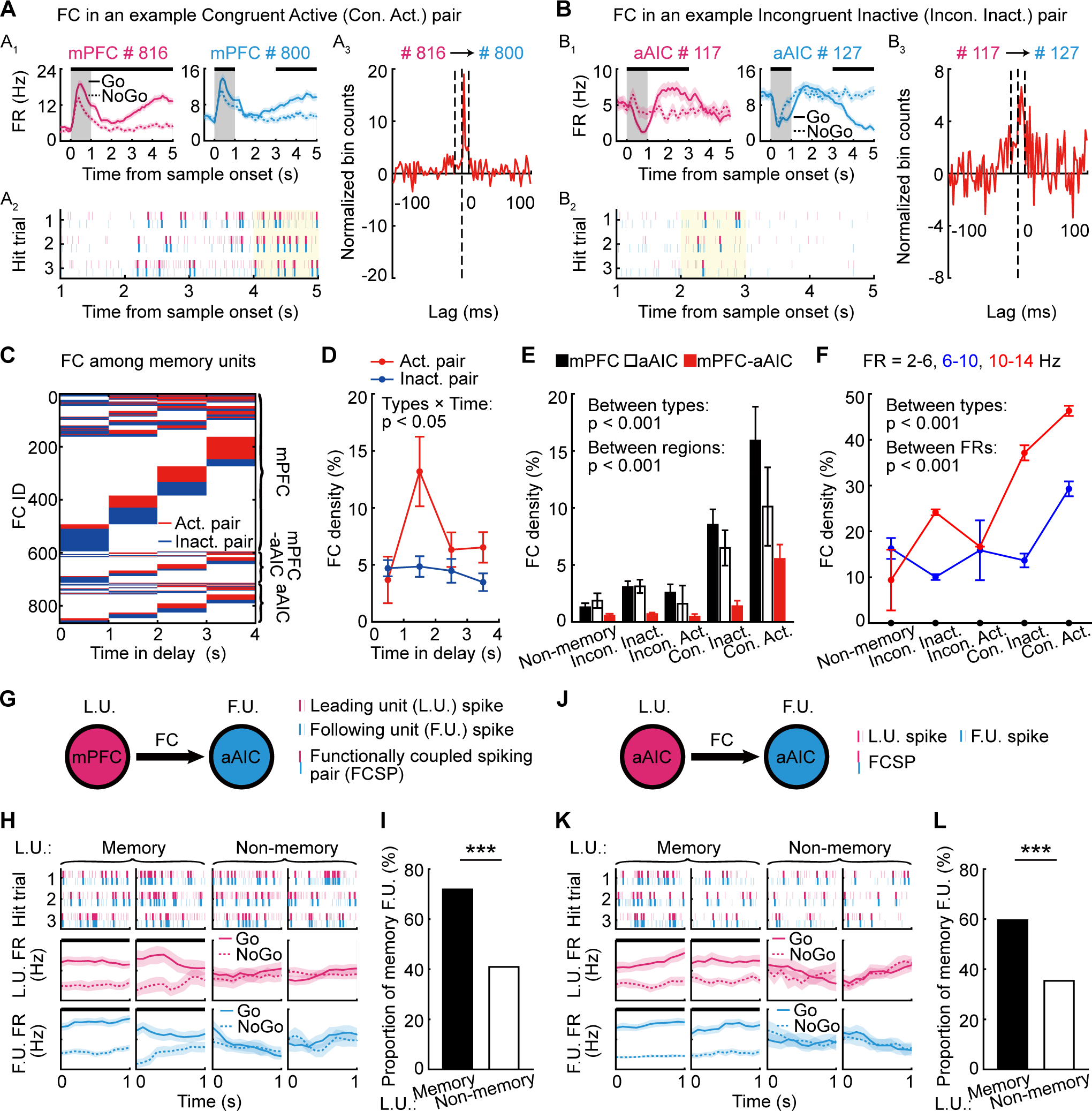
Preferential functional coupling between memory neurons. (A) Functional coupling (FC) of an example congruent active pair (mPFC neuron # 816 and # 800). (A1) PSTH plots of the two example neurons. (A2) Raster plot showing spike timestamps in three example hit trials. Pink and blue bars indicated for spikes of the neuron # 816 and # 800, respectively. Thick line pairs indicated functionally coupled spiking pairs (FCSPs). Yellow region represents the period in which spike trains were used in the spike-correlogram analysis. (A3) Z-scored shift-predictor-subtracted cross-correlogram (SSCC) between spike trains of the two neurons. A prominent and displaced peak on the right side indicated the directional FC from neuron # 816 to # 800. Statistics: p < 0.05, Student *t*-test with Bonferroni correction. Dotted lines represented lag times of -10, 0, and 10 msec (from left to right) after spiking of leading neuron # 816. (B) As (A) for FC of an example incongruent inactive pair. (C) Heatmap showing the temporal profiles of significant FC among memory neurons during the delay period in hit trials. (D) FC densities over the delay period in the hit trial. Statistics: Two-way ANOVA, main effect of types of neuronal pairs F (1,3) = 9.61, p = 2.50 × 10^−3^; main effect of interactions between pair types and time in delay F (3,3) = 3.52, p = 0.018. Error bars represent mean ± SEM. (E) FC density of various types of neuronal pairs. Statistics: Two-way ANOVA, for the comparison between types: F (4,8) = 40.08, p = 3.66 × 10^−26^; between regions: F(2,8) = 31.28, p = 6.64 × 10^−13^. (F) FC density controlling for firing rate (FR). Statistics: Two-way ANOVA, for the comparison between types: F (4,8) = 11.08, p = 4.23 × 10^−6^; between activity levels: F (2,8) = 75.28, p = 4.07 × 10^−14^. (G) Diagram showing preferential FC between memory neurons, from mPFC to aAIC neurons. (H) Example FC neuronal pairs. Raster plots of three example trials (top), PSTH of leading (middle) and following (bottom) neurons. Left two columns show LUs as memory neurons, and right two column show LUs as non-memory neurons. Noted that the following neurons tended to be memory neurons with memory leading neurons. (I) The proportion of following neurons capable of encoding STM information, with memory (black) or non-memory (white) neurons as the leading neuron. Statistics: p = 1.43 × 10^−8^, chi-square test. (J) As (G) within aAIC neurons. (K) As (H) for example FC neuronal pairs in aAIC. (L) As (I) for neuronal pairs with significant FC in aAIC. Statistics: p = 3.57 × 10^−7^, chi-square test.

See also Figure S5.

Neuronal pairs were classified as ‘congruent’ or ‘incongruent’ according to whether two memory neurons preferred the same memory information. Because that many neurons exhibited memory-encoding ability only transiently during the delay period (Figures 2F, S3C, and S3F), we defined neuronal pair to be ‘active’ if both neurons could simultaneously encode memory during a time bin, and ‘inactive’ otherwise. Spike-correlogram analysis for each time bin revealed predominantly transient patterns for FC (examples in Figures 3A2, 3B2, S5A2-S5F2; population in Figure 3C). No difference in FC density could be detected between the active and inactive neuronal pairs in the first second of the delay period. However, much more FC occurred in the active neuronal pairs in the later delay period than in the inactive pairs (Figure 3D).

We observed the highest FC in the congruent and active pairs, and the lowest FC among the non-memory neurons (Figure 3E). Local FC between mPFC neurons was higher than that between aAIC neurons. The local FC within these two regions was higher than cross-region FC, probably due to large distance for cross-region neuronal pairs (Figure 3E). A trivial explanation for the difference in FC density could be the higher firing rate (FR) in congruent active memory neurons (Figure S5G). We therefore controlled the difference in FR by separately calculating FC density for the neurons within a given range of FR (range size: 4 and 6 Hz for analysis in hit and CR trials, respectively). The neuronal pairs with higher FR indeed exhibited higher FC (Figures 3F and S5H). But importantly, the congruent and active pairs still exhibited a higher FC density within similar FR range (Figures 3F and S5H). Therefore, the neurons sharing the same memory selectivity are strongly coupled, even after controlling the FR difference.

Memory neurons could either increase or decrease FR during the delay period comparing to the baseline period (as in Figures 2E, 2J, 2O, and 2P). We observed that FC was predominantly observed in the neurons with increased firing modulation during the delay period (Figure S5I). This is in line with the significant correlation with behavioral performance of these neurons (Figures 2J and 2P).

We wondered whether our simultaneous recording could reveal more subtle FC with memory selectivity. By zooming in to the mPFC-aAIC neuronal pairs with FC, we examined the influence of the coding ability of the leading mPFC neurons to that of the following aAIC neurons. As shown in the example pairs in Figure 3H, after an event of FCSP of a leading mPFC neuron with STM-coding ability, we tended to find that the following aAIC neuron also exhibited STM-coding ability. Overall, the aAIC neurons receiving FC from the mPFC memory neurons were more likely to exhibit memory selectivity than those receiving FC from non-memory mPFC neurons (Figure 3I). Similar patterns were observed for local FC pairs of mPFC and aAIC neurons (Figures 3J-3L and S5J). Therefore, FC analysis revealed preferential functional connection among memory neurons.

### Importance of mPFC-aAIC Projections to DRT Performance

We then utilized pathway-specific optogenetic manipulation to examine its potential causal contribution to behavior. To suppress the activities of mPFC axon terminals in aAIC, AAV-CaMKII_α_-NpHR was injected in mPFC to express NpHR in excitatory neurons and optical fibers were inserted in aAIC (Figures 4A, 4B, and S6D). AAV-CaMKII_α_-YFP was expressed in the control group. Laser illumination (532 nm) was applied during the delay period for all trials throughout the learning phase (top panel of Figure 4C). Optogenetic effects on neuronal activity in aAIC were verified by electrophysiological recordings (Figures S7A and S7B). Optogenetic suppression of mPFC-aAIC pathway during the delay period impaired the CR rate, discriminability (*d’*) and lick efficiency during the learning but not the well-trained phase (comparing NpHR and control groups, Figures 4C, 4D, S6A, and S6B), without any change in the lick rate in the Go trials (Figure 5D) and the proportion of aborted trials (Figure S6C). To further control for the non-memory components of the DRT performance, we further suppressed mPFC-aAIC pathway in the control OGNG task (as in Figure 1I and top panel of Figure 4E) and observed no change to performance (Figure 4E).

**Figure 4.**
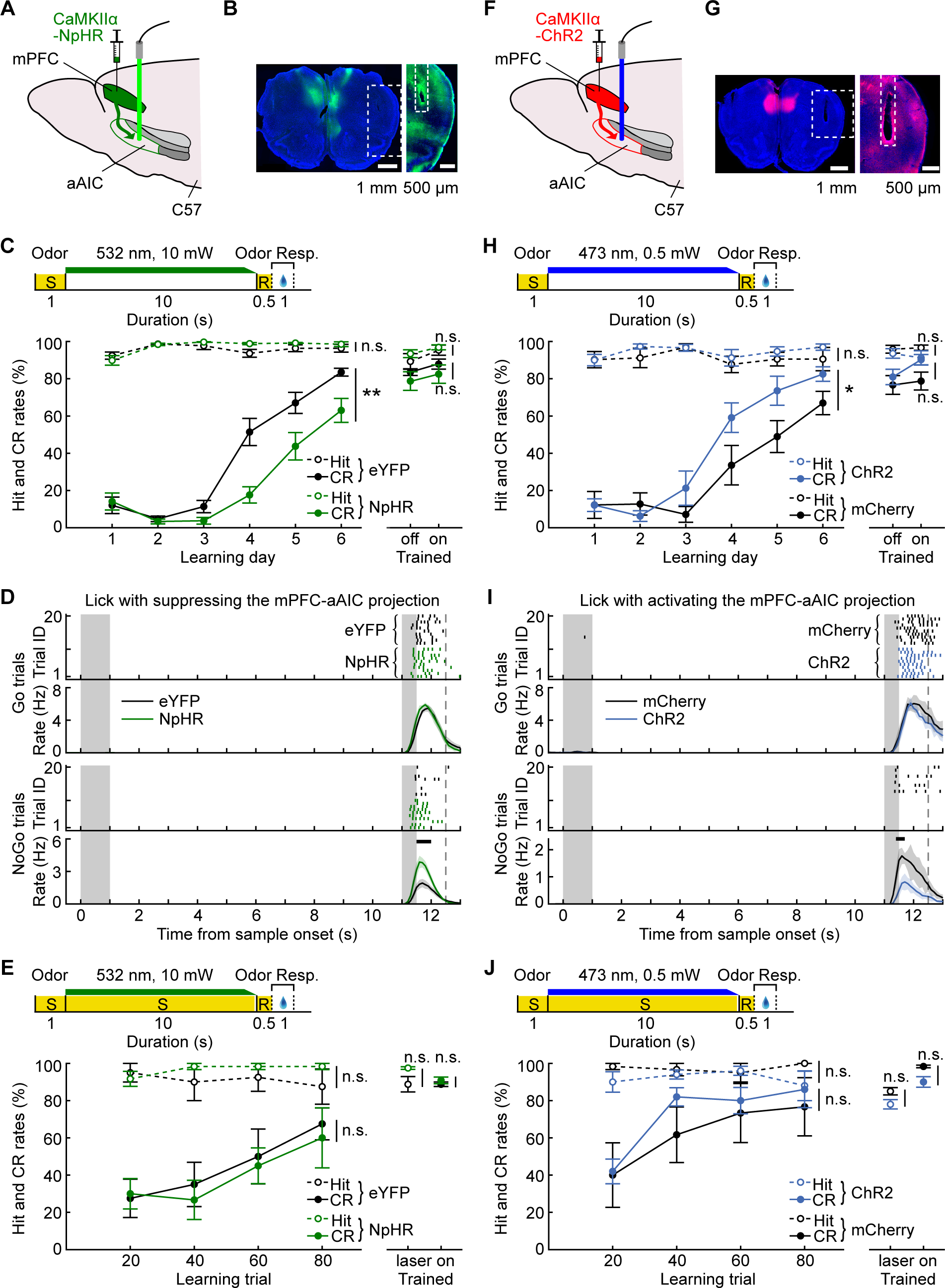
Importance of mPFC-aAIC projections to DRT performance. (A) Schematic diagram for optogenetic suppression of mPFC-aAIC projections. (B) Histology images showing the location of virus CaMKIIα-NpHR in mPFC (left) and optical fibers on top of aAIC (right). (C) Hit and CR rates of mice in performing DRT, for optogenetic suppression of delay-period activity of mPFC-aAIC projections (green) and control (black) groups. Top: schematic of optogenetic suppression. For the hit rate, learning day: F (1,21) = 1.81, p = 0.19; trained: p = 0.45 for laser off/on. For the CR rate, learning day: F (1,21) = 11.47, p = 0.0028; trained: p = 0.22 for laser off/on. Statistics: Tw-ANOVA-md for learning-day performance, n = 12 and 11 mice for NpHR and eYFP groups, respectively; Mann-Whitney U test for the well-trained phase, n = 8 and 7 mice for NpHR and eYFP groups, respectively. Error bars indicate mean ± SEM. (D) Lick raster and lick rate for all mice in Go and NoGo trials. Statistics: p < 0.05, cluster-based permutation test. Shaded area represents mean ± SEM. (E) Hit and CR rates of mice in performing OGNG task, for optogenetic suppression of delay-period activity of mPFC-aAIC projections (green) and control (black) groups. Top: schematics of optogenetic suppression. For the hit rate, learning day: F (1,8) = 0.70, p = 0.43; trained: p = 0.52. For the CR rate, learning day: F (1,8) = 0.11, p = 0.75; trained: p = 0.55. Statistics: Tw-ANOVA-md for learning-day performance, Mann-Whitney U test for the well-trained phase. n = 6 and 4 mice for NpHR and eYFP groups, respectively. (F) As (A) for activation of mPFC-aAIC projections. (G) As (B) for virus CaMKIIα-ChR2. (H) Hit and CR rates of mice in performing DRT, for optogenetic activation of delay-period activity of mPFC-aAIC projections (blue) and control (black) groups. For the hit rate, learning day: F (1,12) = 1.31, p = 0.27; trained: p = 0.39 for laser off/on. For the CR rate, learning day: F (1,12) = 5.98, p = 0.031; trained: p = 0.44 for laser off/on. Statistics: Tw-ANOVA-md for learning-day performance, n = 7 mice for both groups; Mann-Whitney U test for the well-trained phase, n = 7 and 6 mice for ChR2 and mCherry groups, respectively. (I) As (D) for activation of mPFC-aAIC projections. (J) As (E) for activating mPFC-aAIC projections. For the hit rate, learning day: F (1,9) = 1.83, p = 0.21; trained: p = 0.35. For the CR rate, learning day: F (1,9) = 0.34, p = 0.58; trained: p = 0.36. Statistics: Tw-ANOVA-md for learning-day performance, Mann-Whitney U test for the well-trained phase. n = 5 and 6 mice for ChR2 and mCherry groups, respectively.

See also Figure S6.

**Figure 5.**
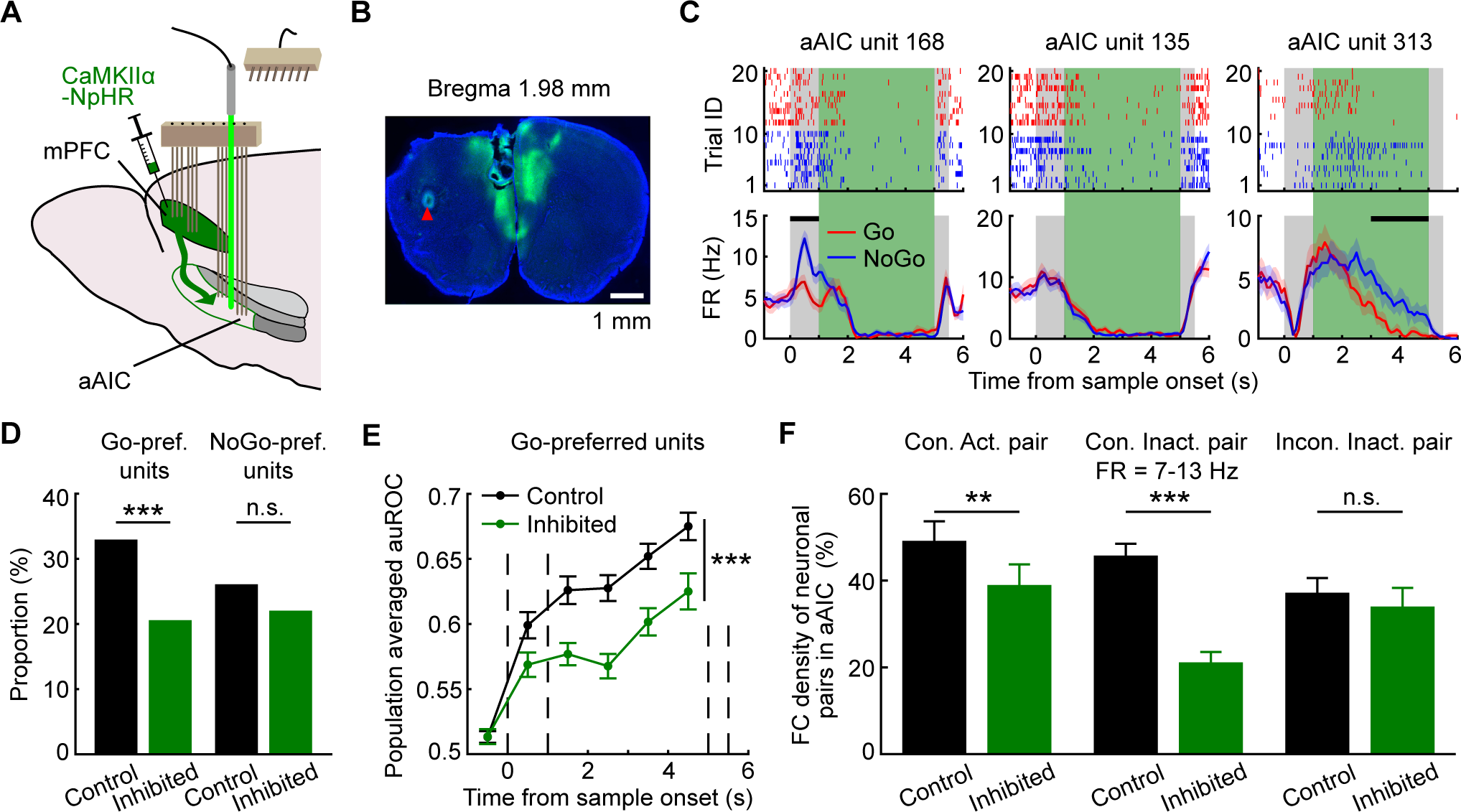
Silencing mPFC-aAIC projections impaired STM-coding ability and FC of memory neurons in aAIC. (A) Diagram for simultaneous optogenetic suppression and single-unit recording. (B) Histology image showing example recording site and virus expression of AAV-CaMKIIα-NpHR. (C) Spike raster (top) and FR (bottom) of three example aAIC neurons, following suppression of mPFC-aAIC projections. Green shadow indicated period of laser application. (D) Proportion of Go-preferred (left, p = 1.19 × 10^−4^) and NoGo-preferred (right, p = 0.19) aAIC neurons, for suppression of mPFC-aAIC projections (green) and control (black) groups. Statistics: chi-square test. (E) Memory-coding ability of Go-preferred aAIC neurons as shown by auROC for sample odors, following suppression of mPFC-aAIC projections. For the comparison between groups: F (1,213) = 19.29, p = 1.77 × 10^−5^. Statistics: Tw-ANOVA-md, n = 144 and 71 neurons for control and suppression groups, respectively. Error bars represent mean ± SEM. (F) FC density of neuronal pairs of memory aAIC neurons controlling for FR, following suppression of mPFC-aAIC projections. Statistics: bootstrap test, p = 0.0020, 0, and 0.096 from left to right. Error bars indicate 95% CI from bootstrap of 1000 times. Noted that congruent neuronal pairs (Con.) showed lower FC density following optogenetic suppression.

See also Figure S7.

Conversely, to examine whether activating mPFC-aAIC projections could enhance STM performance, virus Channelrhodopsin-2 (AAV-CaMKII_α_-ChR2) or control fluorescence protein (CaMKII_α_-mCherry) was expressed in mPFC (Figures 4F, 4G, and S6H). We also verified the effects of optogenetic activation of mPFC-aAIC projections on the neuronal activity in aAIC (Figures S7G and S7H). We found that DRT performance was improved by delay-period laser illumination during the learning but not the well-trained phase (Figures 4H, 4I, S6E, and S6F), without affecting the lick rate in the Go trials (Figure 4I) and with a slight decrease in the proportion of aborted trials (Figure S6G). Activation of mPFC-aAIC pathway in the control OGNG task did not change performance (Figure 4J).

Therefore, optogenetic suppression and activation of the delay-period activity of mPFC-aAIC projections impaired and improved DRT performance, respectively. Thus, the delay-period activity of mPFC-aAIC projections was critical for DRT behavioral performance and memory maintenance during the learning phase.

### Bidirectional Modulation in STM-Coding and FC in aAIC by Prefrontal Projections

To understand the neural mechanisms underlying the behavioral modulation following optogenetic suppression of mPFC-aAIC projections, we recorded aAIC neuronal activity while optogenetically suppressing the activity of mPFC-aAIC projections in mice learning DRT. AAV-CaMKII_α_-NpHR was expressed in mPFC (Figures 5A and 5B; see also Figure S2A) and op-tetrodes were inserted into both mPFC and aAIC, with optical fibers on top of aAIC. We focused our attention on the modulation in the activities of aAIC neurons. Among 345 aAIC neurons recorded from 6 mice (examples in Figure 5C), we observed a significant reduction in the proportion of memory neurons in optogenetic-suppression group, as compared to control mice (Figures 5D, S7D, and S7F). Moreover, suppressing the projections also significantly reduced the coding ability of transient neurons in aAIC during the delay period, as shown by the receiver operating characteristic (ROC) analysis (left panel of Figure S7C). The reduction in both the proportion and coding ability of memory neurons was observed for the Go-preferred neurons, but not for the NoGo-preferred neurons (Figures 5D, 5E, and S7E), in line with the correlation of the Go-preferred neurons with behavioral performance (Figure 2J). The reduction in the proportion of memory neurons was observed in both sustained and transient neurons (Figure S7D). This was different from the olfactory delayed paired association (ODPA) task^18^, in which suppressing the delay-period activity of mPFC-aAIC projections only reduced the proportion of transient neurons.

What’s the local mechanism in aAIC neurons underlying the reduction of information maintenance (Figures 5D, 5E, S7C-S7F) following suppression of mPFC-aAIC pathway? Our analysis revealed a higher proportion of following neurons as memory neurons, if the leading neuron is a memory neuron, both for cross-region FC from mPFC to aAIC (Figures 3G-3I) and within-region aAIC FC (Figures 3J-3L). We thus hypothesized that the suppression of mPFC input to aAIC might reduce the FC among aAIC memory neurons, which could contribute to impaired coding ability for STM information. We thus examined FC among aAIC neurons following optogenetic suppression of mPFC-aAIC pathways. Indeed, we observed a reduction in FC among the congruent memory neurons, but not among the incongruent neurons (Figure 5F). This phenomenon was only observed for the neurons with FR above 7 Hz (Figure 5F). Therefore, the reduced coding ability in aAIC following optogenetic suppression of mPFC-aAIC projections is associated with lower recurrent functional connectivity within aAIC memory neurons encoding the same STM information.

We then examined the changes in aAIC neuronal activity following optogenetic activation of the mPFC-aAIC projection, by expressing AAV-CaMKII_α_-ChR2 in mPFC (Figures 6A and 6B; see also Figure S2A). Laser illumination in aAIC was applied during the delay period for all trials throughout learning the DRT task. Among 377 aAIC single units recorded from 6 mice for the ChR2 group (examples in Figure 6C), optogenetic activation resulted in a significant elevation of the proportion of aAIC memory neurons during the delay period comparing to that of the control mice (Figures 6D and S7L). Activation of mPFC-aAIC projections increased the proportion of the Go-preferred neurons in aAIC, without changing that of the NoGo-preferred neurons (Figure 6D), in line with the results of optogenetic suppression (Figure 5D). The increase in the proportion and coding ability of memory neurons in aAIC was limited to the transient neurons (Figures 6E, S7I, S7J, and S7K). Furthermore, the coding ability of transient memory neurons was significantly enhanced by optogenetic activation of mPFC-aAIC pathway, whereas that of sustained neurons was reduced (Figure S7K). This result further supported the importance of transient neurons in encoding STM information.

**Figure 6.**
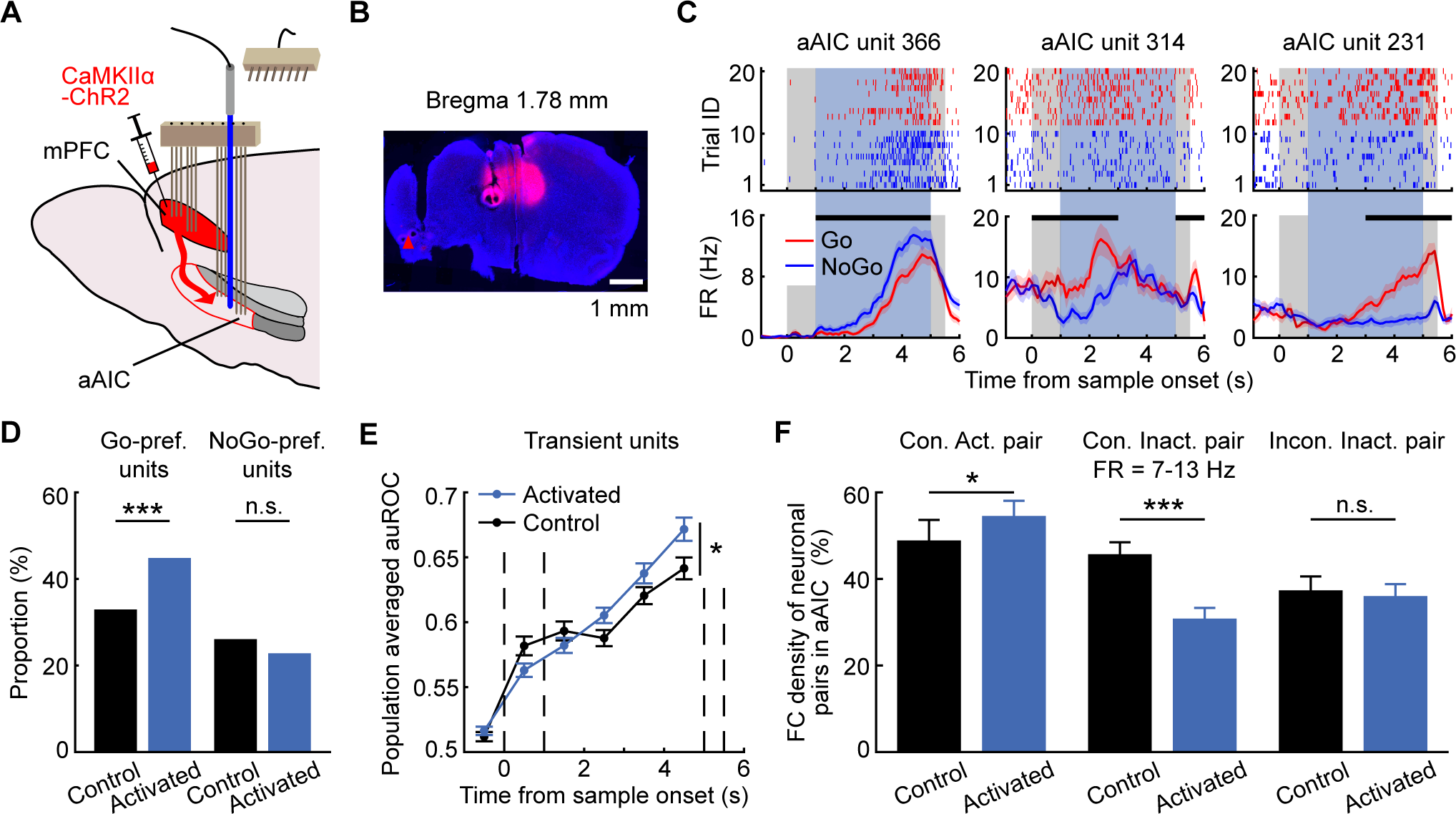
Excitation of mPFC-aAIC projections improved STM-coding ability and FC of memory neurons in aAIC. (A) Diagram for simultaneous optogenetic activation and single-unit recording. (B) Histology image showing example recording site and virus CaMKIIα-ChR2 expression. (C) Spike raster (top) and FR (bottom) of three example aAIC neurons, following excitation of mPFC-aAIC projections. Blue shadow indicated period of laser application. (D) Proportion of Go-preferred (left, p = 5.15 × 10^−4^) and NoGo-preferred (right, p = 0.28) aAIC neurons, for excitation of mPFC-aAIC projections (blue) and control (black) groups. Statistics: chi-square test. (E) Memory-coding ability of transient aAIC neurons as shown by auROC for sample odors, following excitation of mPFC-aAIC projections. For the comparison between groups: F (1,418) = 5.11, p = 0.024. Statistics: Tw-ANOVA-md, n = 208 and 212 neurons for control and excitation groups, respectively. Error bars represent mean ± SEM. (F) FC density of neuronal pairs of memory aAIC neurons controlling for FR, following excitation of mPFC-aAIC projections. Statistics: bootstrap test, p = 0.036, 0 and 0.25 from left to right. Error bars indicate 95% CI from bootstrap of 1000 times. Noted that congruent active neuronal pairs (Con.Act.) showed higher FC density following optogenetic excitation.

See also Figure S7.

We then examined the local mechanism in aAIC neurons underlying the increase of information maintenance (Figures 6D, 6E, S7J, and S7L) following activation of mPFC-aAIC pathway. We observed an enhanced FC among the congruent active memory neurons (Figure 6F). In contrast, the FC density in inactive neurons pairs were reduced following optogenetic activation of the pathway, suggesting that the mPFC-aAIC pathway modulates aAIC memory neurons specifically in the neurons actively encoding the same STM information. Therefore, the enhanced coding ability in aAIC following optogenetic excitation of mPFC-aAIC projections is associated with elevation in recurrent connectivity within aAIC memory neurons.

### Prefrontal Projections Modulate Memory-coding Ability of FC and Efficacy of Information Transfer of Memory aAIC Neurons

Because that the spiking events within FCSP could be important in forming memory-specific cell assemblies, we tested whether the FCSP events could maintain STM information. We firstly examined the coding ability of these FCSP events, based on the FCSP events during the delay period (Figure 7A). Indeed, we found the highest memory-encoding ability for the FCSP events among congruent and active memory neurons (Figure 7B). The proportion of memory-encoding neurons and the density of FC were bi-directionally modulated by optogenetic manipulation in a manner consistent with behavioral modulation (Figures 4C, 4H, 5D, 5F, 6D, and 6F). We wondered whether the coding ability of FCSP could also be modulated in a similar manner. Indeed, we found reduction and improvement of coding ability of FCSP only in congruent active pairs of aAIC neurons following optogenetic suppression and excitation of mPFC-aAIC projections, respectively (Figures 7C and S8A). There was no change in the FR difference between sample odors (Figure S8B), therefore mPFC-aAIC pathways seem not biased toward certain odors in modulation of aAIC coding ability.

**Figure 7.**
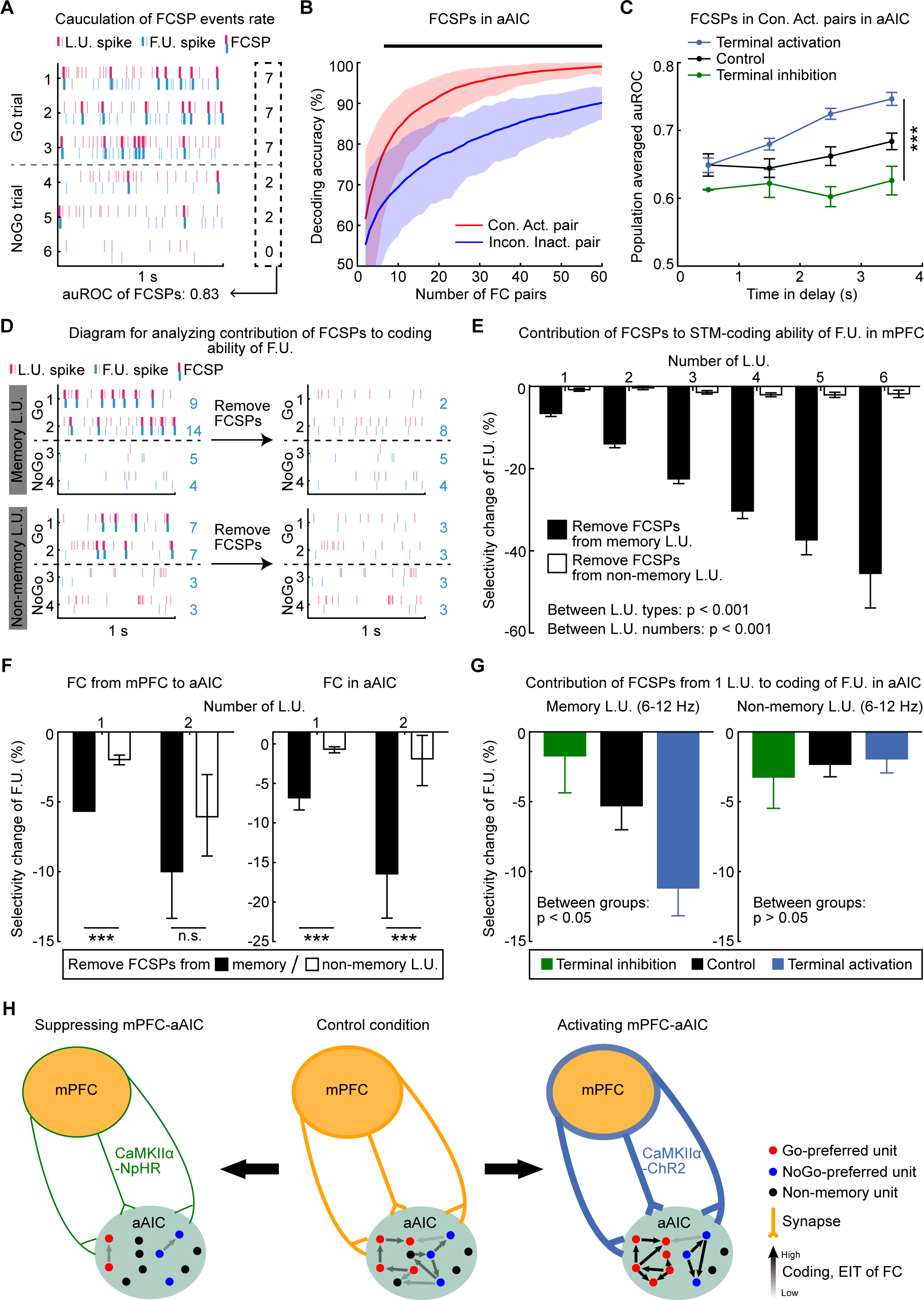
Prefrontal projections modulate memory-coding ability of FC and efficacy of information transfer of aAIC memory neurons. (A) Diagram for calculation of FCSP events rate, indicated by the number on the right side. (B) Decoding accuracy of FCSP events in the congruent active (red) and incongruent inactive (blue) pairs of aAIC neurons. Statistics: p < 0.05, bootstrap test. Shaded area represents 95% CI from bootstrap of 100 times. (C) Memory-coding ability (as shown by auROC for sample odors) of FCSP events in congruent active pairs of aAIC neurons, following optogenetic suppression (green) and activation (blue) of mPFC-aAIC projections, respectively. Statistics: Two-way ANOVA, F (2,6) = 34.43, p = 1.09 × 10^−14^. Error bars represent mean ± SEM. (D) Schematic diagram for contribution of FCSP events to STM-encoding ability of following neurons, with memory (top) or non-memory (bottom) neurons as the leading neuron. (E) Change in memory-coding ability of following mPFC neurons, following the removal of FCSP events from memory (black) or non-memory (white) mPFC neurons. For the comparison between leading-neuron types: F (1,5) = 719.41, p = 9.68 × 10^−141^; between number of leading neurons: F (5,5) = 72.92, p = 8.22 × 10^−72^. Statistics: Two-way ANOVA. (F) As (E) for removing FCSP events across regions (left) and in aAIC (right). Left, p = 0 and 0.069 for 1 and 2 leading neurons with removed FCSP spikes, respectively. Right, p = 0 and 0 for 1 and 2 leading neurons with removed FCSP spikes, respectively. Statistics: bootstrap test. Error bars indicate 95% CI from bootstrap of 1000 times. (G) FCSPs-removal induced change in memory-coding ability of following aAIC neurons with memory (left) or non-memory (right) aAIC neurons as leading neurons (FR controlled), following optogenetic manipulation of mPFC-aAIC projections. Statistics: One-way ANOVA, for memory leading neurons: F (2,151) = 4.15, p = 0.018; for non-memory leading neurons: F (2,82) = 0.24, p = 0.78. (H) Diagram of neural mechanism underlying STM modulation by cross-region projections.

See also Figure S8.

To quantify the contribution of the leading memory neuron to the coding ability of the following neuron, we defined the efficacy of information transfer (EIT) by the magnitude of reduction in the coding ability of the following neurons after removing spikes involved in FCSP. Removing FC spikes from leading neurons with memory encoding ability (Figure 7D) indeed reduced memory selectivity of the following neurons, in a manner correlated with the number of memory-selective leading neurons with removed FCSP spikes (within mPFC, from mPFC to aAIC, and within aAIC in Figures 7E and 7F, respectively). Removing the spikes from non-memory leading neurons did not change the memory encoding ability of the following neurons (Figures 7E and 7F). To control for the possibility that this difference between memory and non-memory leading neurons was due to the difference in their FRs (Figures S8C and S8E), we performed the same analysis for the leading neurons with the same FR ranges and obtained the same results (Figure S8D).

We then examined whether mPFC-aAIC projections contribute to EIT within aAIC neurons. For FC neuronal pairs with both leading and following neurons being memory neurons, EIT was decreased and increased after optogenetic suppression and activation of the mPFC-aAIC projection, respectively (with or without control for FR, Figure 7G, left, S8F). For FC neuronal pairs with non-memory neurons as the leading neurons, projection-specific perturbations didn’t change EIT (Figure 7G, right, S8F). Therefore, the mPFC-aAIC projection bi-directionally modulated the efficacy of information transfer between aAIC memory neurons.

## DISCUSSION

In this study, we have examined the functional role of long-range projections from mPFC to aAIC in modulating STM behavior and the underlying circuitry mechanisms within aAIC. We found that optogenetic suppression and excitation of the delay-period activities of this pathway impaired and improved performance in head-fixed mice learning to perform an olfactory STM task, respectively. Such bidirectional modification in performance was accompanied by consistent changes in the proportion of memory-encoding neurons and memory-specific functional coupling in aAIC. The efficacy of information transfer was also bi-directionally modulated by optogenetic manipulation. Therefore, long-range projections could modulate behavior through providing both information-relevant inputs and driving forces underlying recurrent connectivity among memory neurons encoding the same information (Figure 7H).

Our results of memory-specific FC in mPFC-aAIC circuitry are in line with the cell assembly hypothesis^19–22^, in which sequential flow of neuronal activity across memory-encoding neurons and regions can maintain memory. Importantly, pathway-specific optogenetic perturbation induced consistent modulation in behavioral performance, strength and coding ability of memory-specific FC, as well as efficacy of information transfer as quantified by removing spikes involved in FCSP (Figures 4-7), supporting the causal involvement of correlated activities as detected in FCSPs. The ephemeral correlated activity might help to establish memory-specific circuits in a manner consistent with ‘fire together wire together’ principle, following local learning rule such as spike-timing dependent plasticity^103^. Further studies examining the synaptic strength will be necessary to elucidate the involved synaptic-learning rule in establishing such cell assemblies.

Both sustained^12, 17, 79, 80^ and transient^88, 89, 104–107^ patterns of population activity can readily encode memory-specific information. Here we observed an increased proportion of sustained memory neurons from sensory-oriented to motor-oriented task, with the proportion 2.9% to 7.8% (p < 0.05, chi-square test) for mPFC and 1.1% to 11.4% (p < 0.001, chi-square test) for aAIC (result for sensory-oriented task from ref.^18^). For both sustained and transient patterns of delay-period activity, the temporal patterns governing the activity interaction among STM-encoding neurons were transient in nature (Figure 3C). We also observed strong correlation of transient neurons upon optogenetic manipulation of mPFC-aAIC projections (Figures 6E, S7C, S7D, and S7J). Therefore, both transient and sustained activity patterns could be important for the motor-oriented STM task.

Our results showed that Go-preferred neurons were actively engaged in olfactory motor-oriented STM task. First, the delay-period activity of Go-preferred neurons correlated with behavioral performance (Figures 2J and 2P). Second, the proportion of Go-preferred neurons exhibited consistent modulations with changes in behavioral performance induced by optogenetic perturbations (Figures 5D and 6D). Further physiological or imaging studies with cell-type specificity are required to examine the neuronal types of aAIC neurons receiving mPFC inputs and mPFC neurons projecting to aAIC.

Our FC analysis was based on the results of spike-correlogram, which reflected the correlated firing within 10 msec. A better approach is to use whole-cell recording *in vivo* to examine the post-synaptic potential following presynaptic firing, as performed in studying synaptic inputs from thalamus to somatosensory cortex^108^. Future studies using such a method with resolution of synaptic potential is necessary to definitively elucidate the importance of mPFC-aAIC projections in regulating mono-synaptic strength between aAIC memory neurons.

Long-range connections are cornerstone of inter-area communications and vital for brain functions. Here we uncovered the importance of the long-range projections from mPFC to aAIC in bidirectional modulation of STM behavioral performance, as well as consistent changes in the coding ability of aAIC neurons, the density and coding ability of memory-specific FC, and the efficacy of information transfer within aAIC neurons. Thus, prefrontal projections modulate functional connectivity and coding ability of insular neurons to support short-term memory.

## ACKNOWLEDGMENTS

We thank Dr. Muming Poo for his comments on the manuscript. The work was supported by the Innovations of Science and Technology 2030 from the Ministry of Science and Technology of China (Grant No. 2021ZD0203601), National Key R&D Program of China (Grant No. 2019YFA0709504), National Natural Science Foundation of China (Grant No. 31827803 and 32161133024), the Strategic Priority Research Program of the Chinese Academy of Sciences (Grant No. XDA27010400 and XDB32010100), the Shanghai Municipal Science and Technology Major Project (Grant No. 2018SHZDZX05 and 2021SHZDZX), the Lingang Laboratory (Grant No. LG202105-01-01), Shanghai Pilot Program for Basic Research-Chinese Academy of Science, Shanghai Branch (JCYJ-SHFY-2022-010).

## AUTHOR CONTRIBUTIONS

J.Y. and C.Y.L. designed the study. J.Y. conducted the optogenetics and extracellular recording experiments and analyzed these data. R.Q.H., H.M.F., and J.C.H. contributed to the preparation of virus injection, electrodes construction and histology. Q.C. provided help in analysis and interpretation of extracellular recording data. J.Y. and C.Y.L. interpreted the data and wrote the manuscript.

## STAR*METHODS

### KEY RESOURCES TABLE

2 KEY RESOURCES TABLE
| REAGENT or RESOURCE | SOURCE | IDENTIFIER |
| --- | --- | --- |
| Virus strains |  |  |
| AAV-CaMKIIα-eNpHR3.0-EYFP | Deisseroth lab | Addgene plasmid # 26971;<br>RRID:Addgene_26971 |
| AAV-CaMKIIα-EYFP | OBiO Technology<br>(Shanghai) Corp., Ltd | AOV016 |
| AAV-CaMKIIα-hChR2(H134R)-mCherry | Deisseroth lab | Addgene plasmid # 26975;<br>RRID:Addgene_26975 |
| AAV-CaMKIIα-mCherry | Deisseroth lab | Addgene plasmid # 114469;<br>RRID:Addgene_114469 |
| Experimental models: Organisms/Strains |  |  |
| Mouse: C57BL/6 | Shanghai Laboratory<br>Animal Center<br>(SLAC), CAS, China | RRID:MGI:2159769 |
| Mouse:<br>B6.Cg-Tg(Slc32a1-COP4*H134R/EYFP)8Gfng/J | Jackson lab | RRID:IMSR_JAX:014548 |
| Software and algorithms |  |  |
| ImageJ | National Institutes of<br>Health (NIH) | RRID:SCR_003070 |
| Offline Sorter | Plexon | RRID:SCR_000012 |
| MATLAB | MathWorks, Inc | RRID:SCR_001622 |
| Other |  |  |
| SOLIDWORKS | Dassault Systèmes<br>SOLIDWORKS Corp. | Version 2017<br>https://www.solidworks.com/ |

## RESOURCES AVAILABILITY

### Lead contact

Further information and requests for resources and reagents should be directed to and will be fulfilled by the Lead Contact, Chengyu T. Li.

### Materials availability

This study did not generate new unique reagents.

### Data and code availability

The C codes for behavioral training and matlab codes for data analysis in this study are available from GitHub website: https://github.com/JianYao1992/Motor-oriented-STM

Raw electrophysiology recording files, due to their size (multiple terabytes), are available upon reasonable request.

## Experimental model and subject details

### Mice

In the current study, mice of 14 - 18 weeks of age and 25 - 30 g of weight were used. Wild-type mice were male adult C57BL/6 mice (Shanghai Laboratory Animal Center (SLAC). VGAT-ChR2^+^ mice were B6.Cg-Tg(Slc32a1-COP4*H134R/EYFP)8Gfng/J male mice (Jackson Lab), accompanied by male littermates as controls. Mice were group-housed (4-6/cage) under a 12-h light-dark cycle (light on from 7 a.m. to 7 p.m.). The reported behavioral and electrophysiological results were collected from a total of 55 wild type mice, 29 VGAT-ChR2^+^ mice, and 26 littermates. All animal studies and experimental procedures were performed according to the commonly held animal care standards and have been approved by the Institutional Animal Care and Use Committee of the Center for Excellence in Brain Science and Intelligence Technology, CAS, Shanghai, China.

## Method details

### Behavioral setup

We utilized the olfactory delayed-response task (DRT) and one related control task (olfactory Go/NoGo (OGNG) task) in head-fixed mice (Figures 1A3 and 1F3 for DRT, Figures 1D and 1I for OGNG task) with the olfactometry apparatus enclosed in sound-proof training boxes, as in ref.^37^. An embedded system, custom built around a PIC Digital Signal Controller (dsPIC30F6010A, Microchip, Chandler, AZ), was used to switch on/off the solenoid valves and peristaltic pumps for respectively controlling olfactory cue and water flow in millisecond temporal resolution. Total length of odor-delivery tubes (inner diameter: 2.5 mm; outer diameter: 4.0 mm) was minimized to increase the turnover rate of odorants. All odor reagents were from Sigma-Aldrich, St. Louis, MO. Propyl formate (S1, 245852), 1-butanol (S2, B7906), butyl formate (S3, 261521), ethyl acetate (S4, 270989), 1-pentanol (S5, 398268), 3-methyl-2-buten-1-ol (S6, W364703) and heptane (R, 246654) were kept as stock odorants and evaporated in an air-tight bottle. S1, S2 and R were used for the DRT in the learning phase and suppression of mPFC delay-period activity for the DRT in the well-trained phase. S3, S5, and R were used for suppression of aAIC and suppressing and activating the projection from mPFC to aAIC for the DRT in the well-trained phase. S3, S4, S5, S6, and R were used for the OGNG task in the learning and well-trained phases. The air flow rate was controlled at 2.0 L/min during and between odor deliveries. The odorant vapor was exposed to individually controlled air flow and then mixed with air at the concentrations (v/v) 1:20, 1:8, 1:8, 1:20, 1:4, 1:6.67, 1:20 for S1, S2, S3, S4, S5, S6, and R, respectively. The readout of photoionization detector (200B miniPID, Aurora Scientific Inc, St., Aurora, Canada) for measuring the concentration of odor during the delay period dropped to the baseline level within 0.5 sec after shut-off of the valve, which indicated an efficient clearance of residual odor. About 2.5 - 3.0 _μ_L water was provided in the response window in the hit trials. We detected licking events by a capacitance sensor for optogenetic behavioral experiments, and an infra-red beam break detector for electrophysiological recording. An infra-red camera was placed under the water port for monitoring behavioral states of mice. In order to minimize any potential bias in behavioral results, which were recorded by a computer using customized software written in Java (Oracle, Redwood Shores, CA), intervention from human was not permitted at any time during training.

### Behavioral training

In the DRT, one of two sample odors was presented for 1 sec at the start of a trial, followed by a fixed delay-period for 4 or 10 sec, then a test odor for 0.5 sec. S1 odor signaled a Go trial with water reward if mice liked within response window after the response cue. Conversely, S2 odor signaled a NoGo trial without water reward. Hit and false alarm (FA) was defined as the detection of licking events in the response window in a Go and NoGo trial, respectively. A reward of 2.5 - 3.0 _μ_L water (∼60 - 100 msec in duration) was triggered immediately after a hit response. Mice were neither rewarded nor punished following FA, miss (no lick in a Go trial) or correct rejection (CR, no lick in a NoGo trial) responses. A trial in which licking events were detected during the delay period was aborted immediately until the start of next trial. The results of these aborted trial were not included in analysis. Mice performed 160 trials each training session for optogenetic and electrophysiological experiments. A fixed inter-trial interval (ITI) of 10 sec (for 4-sec delay) or 15 sec (for 10-sec delay) separated consecutive trials. Mice were supplied with free water until satiety after each training session.

The OGNG task was set as a control experiment. Mice were trained to perform the OGNG task in which the sample odor was present during the pseudo ‘delay’ period making up the gap between the sample and test odors in the olfactory DRT. The OGNG task and the DRT shared the same durations of all task events.

Before the behavioral training, mice were water restricted for 1 - 2 days. Water was accessible only during and immediately after training. Care was taken to keep body weight above 80% of pre-training levels. Training included habituation, learning and well-trained phases. In the habituation phase, mice were head-fixed in olfactometry apparatus for 30 minutes and trained to lick water from the water port, encouraged by the automatically delivered water at the beginning. Mice learnt to trigger water rewards more than 300 _μ_L for 2 - 4 minutes typically in 1 - 2 days. The habituation phase lasted for 4 days.

The DRT learning phase was then started from the next day, which was defined as Day 1 in the behavioral analysis reported in all figures. Trial structure was pseudorandomly designed to have two Go and two NoGo trials in each miniblock with four trials. In Day 1, to encourage active licking, water was automatically delivered in the response window for the first 30 Go trials. For the remaining trials, mice learnt to initiate licking for water in the response window of the Go trials.

After the learning phase, for the experiment in which the delay-period activity of mPFC neurons was optogenetically suppressed in the well-trained phase, mice were further trained for several more days to a well-trained level (averaged correct rate for consecutive 60 trials > 80%). In the well-trained phase optogenetic perturbation experiments in which aAIC delay-period activityor the projection from mPFC to aAIC was suppressed, sample odors were changed to unfamiliar ones and mice were trained to reach the well-trained level without optogenetic manipulation. Typically behavioral performance was visualized in either sessions (Figures 1B, 1G, 4C, 4H, S1C-S1I, S6A-S6C, and S6E-S6G) or blocks (Figures 1E, 1J, 4E, and 4J). Hit, miss, FA, and CR rates were defined as follows:

Hit rate = num. hit trials / (num. hit trials + num. miss trials)

Miss rate = num. miss trials / (num. hit trials + num. miss trials)

FA rate = num. FA trials / (num. FA trials + num. CR trials)

CR rate = num. CR trials / (num. FA trials + num. CR trials)

Discriminability (*d’*) was also used to quantify performance, as defined by: *d’ = norminv*(*Hit rate*) *– norminv*(*FA rate*), in which *norminv* was the inverse of the cumulative normal function. Conversion of *Hit* or *FA* rate was applied to avoid plus or minus infinity. In conversion, if *Hit* or *False alarm* rate was equal to 100%, it was set to [1-1/(2*n*)]. If *Hit* or *FA* rate was zero, it was set to 1/(2*n*). Here, *n* equaled to the number of Go or NoGo trials, respectively.

*Lick efficiency* was calculated as:

*Lick efficiency = rewarded licking number /* (*rewarded licking number + unrewarded licking number*)

The *rewarded licking number* referred to the number of licking events in the 0.4-sec time window after the offset of test odor, in which most licking events were detected (Figures 1C, 1H, 4D, and 4I) in the Go trials. The unrewarded licking number referred to the number of licking events in the NoGo trials.

*Early lick* was calculated as:

*Early lick = num. aborted trials /* (*num. aborted tirals + num. finished trials*)

All optogenetic behavioral trainings were in a blind design. For optogenetically suppressing mPFC and aAIC neurons experiments, R. Q. Hou labeled mice and injected virus with ID without revealing the genotype. She did not participate in behavioral and optogenetic experiments or data analysis. The genotype of mice or viruses identity would only be revealed after finishing experiments and statistical analysis of individual mouse.

### Virus preparation

The vectors of optogenetics experiments were obtained from AddGene. Packages of AAV8 viruses were provided by Obio Technology Co. Ltd. (China). Viral titers were 7.0 × 10^12^ particles / mL for AAV-CaMKII_α_-hChR2(H134R)-mCherry, 9.8 × 10^12^ particles / mL for AAV-CaMKII_α_-EYFP, 1.5 × 10^13^ particles / mL for AAV-CaMKII_α_-mCherry. AAV-CaMKII_α_-eNpHR3.0-EYFP virus was provided by Taitool Bioscience Co. Ltd. (China), with the titer of 6.3 × 10^11^ particles / mL.

### Stereotaxic virus injection and optical fiber implantation

Before the surgery, mice were anaesthetized with analgesics (Sodium pentobarbital, 10 mg / mL, 120 mg / Kg b.w.) via intraperitoneal injection. All surgery tools, materials and experimenter-coats were sterilized by autoclaving. Surgery area and materials that cannot undergo autoclaving were sterilized by ultraviolet radiation for more than 20 minutes. Aseptic procedures were applied during surgery. Anesthetized mice were kept on a heat map to maintain normal body temperature. Craniotomies (for virus injection: ∼0.5 mm in diameter; for fiber implantation: ∼1 mm in diameter) were made bilaterally above the targeted area. An injecting pipette was pulled from a glass tube (Sutter glass with filament) to a sharp taper and then the tip was grinded to ∼30 μm in diameter with 45^°^ grinding angle by a micro-grinder. A syringe pump (Harvard apparatus) was used for virus loading and injection at the speed of 1 μL / min and 30 nL / min, respectively. For the optogenetic manipulation of the projection from mPFC to aAIC, virus (0.3 μL in volume) was delivered to each hemisphere with the coordinate

1.98 mm anterior from bregma, 1.21 mm lateral from midline and 1.80 mm ventral from the surface of the brain (with an angle of 26^°^) targeting for prelimbic area (PrL), a part of mPFC. After each injection, to prevent virus spilling over, the pipette was left in tissue for at least 10 minutes before slow withdrawal. Two optical fibers (200 μm in diameter, 0.37 NA, 3 and 4 mm in length for PrL and aAIC) with ceramic ferrule were implanted in bilateral PrL with an angle of 26^°^ or in bilateral aAIC vertically. For PrL, the coordinates of fibers were 1.98 mm anterior from bregma, 1.21 mm lateral from midline and 1.50 mm ventral from the surface of the brain; for aAIC, the coordinates of fibers were 2.52 mm lateral from midline, 1.5 mm posterior from inferior cerebral vein (beneath coronal sutures and between nasal bones and frontal bones) and 1.8 mm ventral from the brain surface. Dental acrylic and cement were mixed and applied to connect skull, plate and optical fibers for structural support. After surgery, mice were recovered under a heat lamp or on a heating pad. Antibiotic drug (Ampicillin sodium, 20 mg / mL, 160 mg / Kg b.w.) was i.p. injected each day for three consecutive days following the surgery.

### Laser illumination designs in optogenetic experiment

An optical patch cable (200 μm in diameter, 0.37 NA) was used to connect the implanted optical fiber (through a ceramic sleeve) to the blue (473 nm) or green (532 nm) diode-pumped solid-state laser (BL473T3-050FC and GL532T3-100FC for the blue and green laser, respectively; SLOC, Shanghai, China). Laser power was programmatically controlled by a PIC microcontroller through analog voltage input. Output laser power was measured with a laser power meter (LP1, Sanwa Electric Instrument Co., Tokyo, Japan) and was adjusted to meet experimental requirement. For the experiments with ChR2 in GABAergic neurons (Figures 1A, 1D, 1F, and 1I), laser (473 nm) power was 2 mW at tips of fibers in brain tissue. For the terminal silencing experiments (Figures 4A and 4B), laser (532 nm) power was 10 mW. For the excitation of mPFC to aAIC terminals, 0.5 mW, 473 nm laser was used (Figures 4F and 4G). For optogenetic experiments in the learning phase, laser illumination was provided in all trials in the learning phase (suppression of mPFC or aAIC, day 1-5; perturbation of mPFC to aAIC terminals, day 1-6). This design meant to exert maximal optogenetic effect, which could be compared across groups of mice with different genotypes or different viruses expressed, maximally utilizing the advantage of blind design that we used. For the optogenetic experiments in the well-trained phase, the optogenetic manipulation was arranged in an interleaved block-based laser off/on mode, and each block (total number: 8) consisted of 20 trials; each session started with a laser-off block except for the terminal activation experiment. For the optogenetic experiments in the control OGNG task, laser illumination was provided in all trials in the learning and well-trained phases except that suppressing mPFC neurons in the well-trained phase was arranged in an interleaved block-based laser off/on mode (20 trials / block, total 8 blocks / session) with the first block without laser illumination.

### Immunostaining and imaging experiments

Mice were deeply anesthetized with sodium pentobarbital (120 mg / Kg b.w.) and then perfused transcardially with 20 mL saline followed by 20 mL 4 % paraformaldehyde (PFA, in 0.01 M phosphate buffer saline, PBS, pH 7.4). The brains were removed and kept in 4% PFA at 4℃C overnight, then transferred to PBS. Coronal slices (80 μm in thickness) were obtained using a vibratome and collected in PBS. Slices were incubated with DAPI (C1002, Beyotime, 1:1000 diluted in PBS) for 10-15 min, then mounted and coverslipped after washing three times by PBS (10 min each time). Fluorescence images were then obtained with Olympus VS120 under a 10X (0.45 NA) lens and were further analyzed with ImageJ software (NIH, US).

### Generation of overlaid histology maps

We focused on two coronal brain sections (Bregma +2.10 and 1.94 mm, following the atlas of Paxinos & Franklin 2008) to report optical fibers implantation locations from VGAT-ChR2^+^ mice and then traced out fibers locations on these selected brain slices to obtain area of interests. For each mouse with virus CaMKII_α_-NpHR or CaMKII_α_-ChR2 expressed, viral expression areas on three coronal brain slices (Bregma +2.34, 2.10, 1.94, 1.70, and 1.54 mm) were marked in the matrix figure as area of interests. We summed these figures from different mice (number: n) to generate an overlaid image with number from zero to n, with each successful fluorescence or optical-fiber marking adding one to the figure. Darkness of this summed matrix figure was used to generate the overlaid histology map for silencing mPFC (Figure S1J) or aAIC (Figure S1K) group, optogenetic behavior group of terminal suppression (Figure S6D) or activation (Figure S6H), and electrophysiological group (Figure S2A). The darker pixel indicated the larger number of mice with fibers implanted or virus expressed. Colored dots in Figure S2A indicated lesion sites from mice for electrophysiological experiments including data from mice of control group and same-colored dots marked data from one individual mouse.

### Electrodes array construction and implantation

Multi-electrode array used in extracellular recording were custom-made as described previously^6, 18^. Tetrode was constructed with polyimide-insulated Ni-Chrome wire (12.5 μm core diameter). Wire tips were soldered on printed circuit boards which were connected to preamplifier adaptor. One reference electrode and one ground were set for all electrodes within a drive. Each microdrive was composed of 16 tetrodes with 64 recording channels. For recording activities of mPFC and aAIC neurons simultaneously, chamber was designed with SOLIDWORKS software to target these two brain regions accurately, with 8 tetrodes in each region of the left hemisphere. For two mice with recording only in mPFC, each hemisphere was allocated with 8 tetrodes. Optical fibers were placed adjacent to the electrodes (within 500 μm) under conditions of optogenetic perturbations of neuronal activity. The procedure for implantation of the microdrive was similar to that for implantation of optical fibers.

A microdrive was sterilized by ultraviolet radiation for more than 20 minutes. Mice were deeply anesthetized with analgesics (Sodium pentobarbital, 10 mg / mL, 120 mg / Kg b.w.) before surgery. The center of the electrode array was targeted to 1.98 mm anterior from the bregma, 0.40 mm lateral from the midline, and 1.65 mm ventral from the surface of the brain for mPFC; 1.5∼1.9 mm posterior from the inferior cerebral vein, 2.4 mm lateral from the midline, and ∼2.25 mm ventral from the brain surface for aAIC. Dura mater was carefully removed with surgery needles with less bleeding for the insertion of electrode array. Tissue gel (3M, US) and dental cement were carefully applied without motion between a microdrive and brain tissue. Antibiotic drug (ampicillin sodium, 20 mg / mL, 160 mg / Kg b.w.) was i.p. injected for following three consecutive days after surgery.

### Electrophysiological recording

Before behavioral training, mice were allowed to recover for at least two weeks after the implantation of the electrode array. Neuronal activities were then recorded with a Multichannel Acquisition Processor (Plexon Inc, Dallas, Texas). The recording began after the end of habituation period. Wide band signals (0.5-8000 Hz) from all tetrodes were amplified (× 20000) and digitized at 40 KHz and all data were saved to hard disks. After each training session, the screw was turned anticlockwise for about 90 degree lowering the recording tetrodes for ∼50 μm, to decrease the possibility of resampling the same neurons across different days. Spike sorting was performed based on the spike waveforms by using Offline Sorter software (Plexon Inc). Raw signals were filtered in 250 - 8000 Hz to remove field potentials. Typically signals larger than negative five times the standard deviation recorded at any lead of the tetrode were treated as spike events. Principal component analysis (PCA) was performed for tetrode-waveforms to extract the first three principal components (PCs) that explained the largest variance. Then, ‘Valley-seek’ clustering techniques provided by Offline Sorter were performed in 3D feature space of waveforms (Figure S2B). The signal-to-noise ratio (SNR, the mean peak amplitude of all spikes divided by the standard deviation during baseline, see Figure S2E, S2F, and S2G), false alarm rate (FA, spikes within 2 msec refractory period divided by the total spikes number, see Figure S2C, S2F, and S2G), and averaged firing rate (FR) were measured for controlling the quality of sorted single units. Only clusters with FA less than 0.15% and FR higher than 2 Hz were considered to be single unit with good quality. Neurons with discontinuous spiking activity (stability visualized in the ‘Clusters vs Time’ feature space) during recording were discarded (Figure S2D). For the simultaneously recorded neurons, the cross-correlograms of inter-spike-interval were carefully examined to make sure that good clusters were not contaminated or over-split. After finishing the well-trained phase recording, all recording sites were marked by passing current (50 μA, 1 sec, 0.5 Hz × 10 pulses) through the electrodes, and were verified with immunostaining one day after the electrical lesion (Figure S2A).

### Spike count and selectivity analyses

We analyzed all data by codes written in MATLAB (Mathworks, Natick, MA). For the current research, all neurons were recorded in the learning phase. 816 mPFC neurons and 437 aAIC neurons were recorded from 48 recording sessions (8 mice) for control group; 265 mPFC neurons and 345 aAIC neurons from 36 recording sessions (6 mice) were recorded with optogenetic suppression of the mPFC-aAIC projection; 241 mPFC neurons and 377 aAIC neurons from 36 recording sessions (6 mice) with optogenetic activation of the mPFC-aAIC projection. Averaged firing rates of individual neurons in all trials for different olfactory samples were binned into 100-msec periods, illustrated in peri-stimulus time histogram (PSTH) plots (Figures 2C, 2D, 2M, 2N, 3A1, 3B1, 3H, 3K, 5C, 6C, S3A, S3B, S3I2-S3K2, S4A-S4D, S5A1-S5F1, S7A, S7B, S7G, and S7H). The mean firing rate and standard deviation (SD) in the baseline period (2 sec before the onset of sample odor) were used to convert the averaged firing rate of each time bins into Z-score. Neurons were sorted based on the difference of delay-period averaged firing rate following different sample odors. The Z-scored firing rate was plotted as heatmap using ‘Jet’ colormap defined in MATLAB (Figures 2E and 2O).

For the analysis of selectivity, a neuron was deemed with significant selectivity if the firing rate within any of the 1-sec bin during the delay period (totally 4 bins) was significantly different between two sample odors (based on the Mann-Whitney *U* test with Bonferroni correction). Neurons showing significant selectivity for the entire delay duration were defined as sustained neurons, otherwise transient neurons. For the definition of transient neurons, the selectivity during the significant time bins was required to be significantly different from that of the non-significant time bins. The firing rate differences between any time bin (from t_1_ to t_4_) for each trial were calculated, and the N_trial_number_ × t_n_ × t_n_ difference matrix was subjected to Mann-Whitney *U* test for the two-trial groups with different sample odors. We permutated the time labels and tested the significance of the resultant 2-D matrix of Z-statistics with permutation test of 2000 times (Figures S3A and S3B). We computed neuronal selectivity as the formula: Selectivity = (FR_Go_ – FR_NoGo_) / (FR_Go_ + FR_NoGo_), which resulted in a [-1 1] range.

### Population trajectories

We modified dimensionality reduction analysis^88, 109^ to represent population trajectories (Figure 2H). One dimension in an abstract space of population activity dynamics was defined by averaged firing rate of each mPFC or aAIC neuron for particular sample odor. Neurons recorded across mice and learning days were grouped together to generate high dimensional neural firing patterns. For visualization of this high dimensional data, principal component analysis (PCA) was performed and firing dynamics were projected onto the first three PCs explaining the largest variance. All recorded neurons in mPFC (n=816) or aAIC (n=437) were included to calculate PCA. The first three PCs represented 80% and 83% of variance for mPFC and aAIC, respectively. Binned firing rates (bin size: 100 msec) for all mPFC or aAIC neurons were used as inputs for PCA. The resulting first three PCs explaining the largest variance were used to generate trajectories shown in Figure 2H. The first 20 PCs (representing over 95% variance for both mPFC and aAIC) were used for calculation of distance of population trajectory evoked by different sample odors. We calculated the trajectories by the Euclidian distance of these 20 PCs for each time bin. We used the averaged firing rates of randomly selected neurons in mPFC (n = 600) or aAIC (n = 300) to calculate PCA and generate distance between different sample odors for each run of the distance analysis (total 100 times). To test the statistical significance of the trajectory distance, we performed cluster-based non-parametric permutation test after repeating the shuffle procedure for 1000 times by randomly re-assigning the trial labels. Distance was averaged and then plotted to generate Figure 2H for aAIC (result for mPFC not shown), respectively.

### Cross-temporal decoding analysis

We performed the cross-temporal decoding (CTD) analysis^85, 86^ by using the maximum correlation coefficient (MCC) classifiers to measure the persistence of coding abilities for neurons in mPFC and aAIC. To prevent the potential bias induced by the difference in the number of neurons included in the decoding analysis, we chose the same number of neurons for each resample run with 60 trials randomly selected for each sample stimulus and each selected neuron [for sampling from all and transient neurons, n = 150 and 300 neurons for aAIC (Figures 2G and right panel of S3G) and mPFC (Figure S3D, right), respectively; for sampling from sustained neurons, n = 40 and 54 neurons for aAIC (Figure S3G, left) and mPFC (Figure S3D, left), respectively, due to relatively small proportion]. For each neuron in each cross-validation, 59 trials for each stimulus were used as the training data and the remaining trial as the test data. The training and test data for each stimulus were Z-scored, by mean firing rate and SD of each neuron within each non-overlapping time bin (bin size: 100 msec) calculated from concatenation of training data for each stimulus. Z-scored firing rate of all selected neurons were concatenated together for the conditions ‘Template-Go’, ‘Template-NoGo’, ‘Test-Go’, and ‘Test-NoGo’. Classifiers were systematically trained with different time bins, which were then used to test the data for all the time bins, based on the maximal correlation coefficient of MCC classifier. The truth score of 1 and 0 was assigned if the classification was correct and incorrect, respectively. We produced a 2-D cross-temporal decoding matrix by repeating this process for each time bin. The classification accuracy was estimated over the 60 cross-validation trials for each resample run. This decoding analysis was repeated for 100 times. We reported the averaged results in Figures 2G, S3D, and S3G. We performed the shuffled decoding procedures for 1000 times by randomly re-assigning the IDs of sample odors, to determine the chance level with cluster-based non-parametric permutation test. For the analyses (for aAIC: Figures 2G and S3G; for mPFC: Figure S3D), all neurons, sustained neurons, and transient neurons were pooled as the pseudo populations.

### Spike-correlogram analysis

We adapted the cross-correlogram method from ref.^27, 28^ to measure the directional functional coupling (FC) between simultaneously recorded neurons. For each neuronal pair, raw cross-correlograms were constructed for lag times within 100 msec (2-msec resolution) using spikes recorded during each time bin (bin size: 1 sec). Cross-correlogram for Hit or CR trials was accepted only if number of available spikes exceeded 1000. To control for the shared variance, we adopted a shift-predictor subtracting method: shift predictor was calculated using a one-trial-shifted spike trains and smoothed with moving average of five 2-msec bins, which was then subtracted from raw cross-correlogram resulting in shift predictor-subtracted cross-correlogram (SSCC). We normalized SSCC peak height by SD of the shift predictor to calculate the Z-scored peak height (Figures 3A3, 3B3, and S5A3-S5F3). Significant FC was determined with a p-value threshold of 0.05 (one-sided Student *t*-test, detected within the 10-msec lags, corrected for multiple comparisons). Correlated spike counts within ± 2-msec lags were excluded in our analyses to control for SSCC peak due to common inputs. To test the directionality of FC, which was reflected in SSCC with at least one significant peak, the asymmetry index (AI) of the peak structure for each SSCC was calculated as: *AI =* (*L - R*) */* (*L + R*), in which L and R was the significant SSCC spike counts on the left and right sides of SSCC within the lag time of 10 msec, respectively. A significant SSCC peak with AI > 0 was regarded as a directional pairs with FC. This procedure was repeated for all 7 non-overlapping time bins spanning the whole trial (from 1 sec before the onset of the sample odor to 0.5 sec after the offset of the test odor).

### Functional coupling density analysis

FC density was the proportion of neuronal pairs with FC among simultaneously recorded neurons in all possible pairs. For the analysis of FC density, the cross-correlogram analysis was performed in the correct (Hit or CR) trials. For comparison of the FC densities in different types of neuronal pairs, the switched neurons (showing changed coding preference during delay) were excluded in the analysis (Figures 3E, 3F, 5F, 6F, S5H, and S5I). We classified neuronal pairs of each time bin during delay period into the congruent active, congruent inactive, incongruent active, incongruent inactive, and non-memory pairs, based on the coding preferences for STM information during delay and significant coding abilities in current bin of the two neurons. Neuronal pairs were either organized in sessions (Figure 3E) or summed across all sessions (Figures 3D, 3F, 5F, 6F, S5H, and S5I). For calculation of FC density in the CR trials, the neuronal pairs with the number of CR trials less than 60 were excluded to control for the potential bias of the trial number on the significance of SSCC peak height (Figure S5H).

### Receiver operating characteristics analysis for neuronal spiking activity

The receiver operating characteristic (ROC) analysis was performed to measure the discrimination ability for sample odors of single neurons in the DRT, as in ref.^110^. For each trial, a ‘decision variable’ (DV) was calculated as: *DV = FR_Go(i)_* (*mean*(*FR_Go(j_*_≠*i)*_) *– mean*(*FR_NoGo_*)) for the Go trials and *DV = FR_NoGo(i)_* (*mean*(*FR_Go_*) *– mean*(*FR_NoGo(j_*_≠*i)*_)) for the NoGo trials, in which *FR_Go(i)_* was the PSTH of the i-th Go trial, *mean*(*FR_NoGo_*) was the mean PSTH of all NoGo trials, *mean*(*FR_go(j_*_≠*i)*_) was the mean PSTH of all Go trials without incorporation of the i-th trial. Same denotations applied for the *FR_NoGo(i)_*, *mean*(*FR_Go_*) and *mean*(*FR_NoGo(j_*_≠*i)*_). More similarity to Go trials was represented by larger DV. We calculated auROC for neuronal spiking activity with 1-sec non-overlapping time bins (Figures 5E, 6E, S3I-S3L, S7C, S7E, S7I, and S7K).

### Receiver operating characteristics analysis for events of functionally coupled spiking pair

We treated functionally coupled spiking pair (FCSP) as a computing unit and also calculated auROC of these FCSP. Firing rate of above formula was replaced with FCSP (Figures 7A, 7C, and S8A).

## QUANTIFICATION AND STATISTICAL ANALYSIS

Statistical analyses were performed using MATLAB and presented along with p values in the Figure legends. Statistical significance was defined as p < 0.05 unless stated otherwise.

1) For behavioral data except lick rate, Two-way ANOVA in mixed designs (Tw-ANOVA-md) was used in the learning phase (left panels in Figures 1B, 1E, 1G, 1J, 4C, 4E, 4H, 4J, S1C-S1H, S6A-S6C, and S6E-S6G) and non-parametric Mann-Whitney *U* test was used in the well-trained phase (Figure S1I and right panels in Figures 1B, 1E, 1G, 1J, 4C, 4E, 4H, 4J, S1C-S1H, S6A-S6C, and S6E-S6G). For paired data, Wilcoxon signed-rank test was used (Figures S3E and S3H).
2) Selectivity of single neurons was tested with the non-parametric Mann-Whitney *U* test corrected by Bonferroni method for multiple comparison (Figures 2C, 2D, 2F, 2M, 2N, 3A1, 3B1, 3H, 3K, 5C, 6C, S3A, S3B, S4A-S4D, and S5A1-S5F1). Chi-square test was used to compare the proportions of neurons encoding two samples (without Bonferroni correction: Figures 3I, 3L, 5D, 6D, S3C, S3F, S5J, S7D, and S7J; with Bonferroni correction: Figures S7F and S7L). The coding ability of neurons following pathway-specific optogenetic manipulation was tested with Tw-ANOVA-md (Figures 5E, 6E, S7C, S7E, S7I, and S7K). The coding ability of FCSPs and difference in FRs between different sample odors in FC neuronal pairs with projection-specific optogenetic perturbation were tested using Two-way ANOVA (Figures 7C, S8A, and S8B).
3) For the detection of FC within a pair of neurons, Student *t*-test with Bonferroni correction was used (Figures 3A3, 3B3, 3C and S5A3-S5F3). Significance of comparison for FC densities between types of neuronal pairs (Figures 3D-3F, S5H, and S5I) and contribution of FCSPs in mPFC to coding ability of following neurons (Figures 7E and S8D) were tested with Two-way ANOVA. Significant effect of projection-specific manipulation on contribution of FCSPs in aAIC to coding ability of following neurons and activities of leading neurons was tested using One-way ANOVA (Figures 7G and S8F).
4) We used non-parametric permutation test to examine the difference between real and permutated results (Figures S3A and S3B). Cluster-based non-parametric permutation test^91, 111^ was used to verify the significance of all time-resolved results (Figures 1C, 1H, 2G, 2I, 2J, 2P, 4D, 4I, S3D, and S3G). The inherent correlation in consecutive time points is considered by this method indicating that the true significant time points should be clustered in consecutive time points while the false-positive point is shown in isolated event. Taking the decoding analysis as the example, decoding accuracy of 1000 permutations with randomly shuffled sample odors was calculated. The time bin exceeding the 95^th^ percentile of 1000 times permutation distribution was significant for each permutation. We defined clusters as consecutive significant time bins and constructed the null distribution of maximum cluster test statistics with the maximum cluster size of each permutation procedure. The cluster in real decoding results was regarded as a significant cluster if the size of the cluster exceeded the 95^th^ percentile of the null distribution of the maximum cluster test statistic. Bootstrap test was used in the resample procedures reported in Figures 5F, 6F, 7B, 7F, S8C, and S8E.

**Figure S1. Information associated with optogenetic suppression. Related to Figure 1.**

(A) Waveforms (first row), raster plots (second row), and PSTH plots (third row) of two example mPFC neurons with different modulations by laser.

(B) The proportion (left) and population averaged normalized FR (right) of mPFC neurons with different modulations.

(C) Performance in *d’* of mice in performing DRT, for optogenetic suppression of mPFC delay-period activity (blue) and control (black) groups. Statistics: F (1,19) = 6.07, p = 0.023, Tw-ANOVA-md for learning-day performance, n = 11 and 10 mice for VGAT-ChR2^+^ and VGAT-ChR2^−^ groups, respectively; p = 0.0019 for laser off/on in the well-trained phase, Mann-Whitney *U* test, n = 8 mice for both groups. Error bars indicate mean ± SEM.

(D) Lick efficiency, following suppression of mPFC. Statistics: F (1,19) = 4.75, p = 0.042, Tw-ANOVA-md for learning-day performance; p = 0.0030 for laser off/on in the well-trained phase, Mann-Whitney *U* test. Number of mice as in (C).

(E) Proportion of aborted (early lick) trials, following suppression of mPFC. Statistics: F (1,19) = 1.13, p = 0.30, Tw-ANOVA-md for learning-day performance; p = 0.12 for laser off/on in the well-trained phase, Mann-Whitney *U* test. Number of mice as in (C).

(F) As (C) for suppressing aAIC delay-period activity. Statistics: F (1,13) = 15.77, p = 0.0016, Tw-ANOVA-md for learning-day performance, n = 7 and 8 mice for VGAT-ChR2^+^ and VGAT-ChR2^−^ groups, respectively; p = 0.86 for laser off/on in the well-trained phase, Mann-Whitney *U* test, n = 8 mice for both groups.

(G) As (D) for suppression of aAIC. Statistics: F (1,13) = 14.59, p = 0.0021, Tw-ANOVA-md for learning-day performance; p = 0.57 for laser off/on in the well-trained phase, Mann-Whitney *U* test. Number of mice as in (F).

(H) As (E) for suppressing aAIC. Statistics: F (1,13) = 4.28, p = 0.059, Tw-ANOVA-md for learning-day performance; p = 0.46 for laser off/on in the well-trained phase, Mann-Whitney *U* test. Number of mice as in (F).

(I) Hit and CR rates of mice in performing DRT in the well-trained phase, following suppression of mPFC (blue) or aAIC (orange) delay-period activity. Statistics: Mann-Whitney *U* test, p = 0.49 and 0.0016 for the hit and CR rate, respectively. n = 8 mice for suppressing mPFC and aAIC groups, respectively.

(J) Accumulated histology images showing locations of optical fibers on top of mPFC in mice in Figures 1B and 1E.

(K) Accumulated histology images showing locations of optical fibers on top of aAIC in mice in Figures 1G and 1J.

**Figure S2. Spiking sorting. Related to Figure 2.**

(A) Accumulated images showing unilateral extracellular recording sites in aAIC based on electricity lesion after recording for Figures 2, 5, and 6, with the dots of different color indicated for recording site from different mice. Shaded area indicated accumulated virus expression in mPFC in Figures 5 and 6.

(B) Three example clusters of spikes in the feature space based on PCA of all spikes from one tetrode.

(C) Spike autocorrelogram and cross-correlogram for clusters shown in (B).

(D) Distribution of the same clusters in (B) cross time, showing the stability of the recorded neuron. White arrows in Neuron 2 indicated unstable examples. This type of unstable neurons were excluded from further analysis.

(E) Waveforms of two qualified neurons included for further analysis in (B). Signal to noise ratio (SNR): mean peak amplitude of all spikes divided by standard deviation during baseline.

(F) Autocorrelograms for example neurons. False alarm (FA) rate is the ratio of spikes within inter-spike-interval less than 2 msec. Black for mPFC and blue for aAIC neurons.

(G) Scatter plots of SNR and FA rate of all recorded neurons.

**Figure S3. Transient and sustained neurons in mPFC and aAIC in DRT. Related to Figure 2.**

(A) An example unclassified neuron exhibiting ‘sustained’-like selectivity. Left, PSTH plot. Statistics: p < 0.05, Mann-Whitney *U* test with Bonferroni correction. Right, Z-statistic of FR difference matrix during delay. Statistics: permutation test, with p values indicated by numbers. Because of the large P value between the activity of different time bins, these neurons (n = 7/4 neurons for mPFC/aAIC) were not classified into sustained or transient group.

(B) An example un-classified ‘transient’-like neuron. Note these neurons (n = 30/14 for mPFC/aAIC) were not classified into transient group.

(C) Proportion of sustained (7.84%) and transient (37.87%) mPFC neurons. Statistics: p = 0, chi-square test.

(D) CTD results with sustained (left) and transient (right) mPFC neurons, as Figure 2G.

(E) Persistence in memory-coding ability of transient and sustained mPFC neurons, indicating the stability of information maintenance. Dots indicated for coding persistence of different training time point, which was calculated as the duration of significant decoding from the onset of sample odor to the offset of test odor. Statistics: p = 2.89 × 10^−10^, Wilcoxon signed-rank test.

(F) Proportion of sustained (11.44%) and transient (47.60%) aAIC neurons. Statistics: p = 0, chi-square test.

(G) Same as (D) for sustained (left) and transient (right) aAIC neurons.

(H) Same as (E) for transient and sustained aAIC neurons. Statistics: p = 4.87 × 10^−10^, Wilcoxon signed-rank test.

(I) Cross-trial FR of an example sustained neuron, with auROC value 0.74. (I1) Fano Factor in preferred Go trials. (I2) FR of each single trials. (I3) Distribution of FR of all trials.

(J) As (I) for another sustained neuron, with auROC value 0.89.

(K) As (I) for another sustained neuron, with auROC value 0.99.

(L) Distribution of memory-coding ability of all sustained neurons (n = 114 neurons).

**Figure S4. Selectivity of memory neurons during delay and action periods. Related to Figure 2.**

(A) FR of example Go-preferred neurons, losing selectivity for hit and CR trials after delay.

(B) FR of example Go-preferred neurons, switching selectivity for hit and CR trials after delay.

(C) As (A) for NoGo-preferred neurons.

(D) As (B) for NoGo-preferred neurons.

(E) Proportion of neurons keeping, losing, and switching selectivity in the Go-preferred (left) and NoGo-preferred (right) mPFC neurons.

(F) As (E) for aAIC.

**Figure S5. Preferential functional coupling between memory neurons. Related to Figure 3.**

(A) FC of an example congruent active pair (mPFC neuron # 116 and aAIC # 128). (A1) PSTH plots of the two example neurons. (A2) Raster plot showing spike timestamps in three example hit trials. Pink and blue bars indicated for spikes of mPFC neuron # 116 and aAIC # 128, respectively. Thick line pairs indicated FCSPs. Yellow region represents the period in which spike trains were used in the cross-correlogram analysis. (A3) Z-scored SSCC between spike trains of the two neurons. A prominent displaced peak on the right side indicated the directional FC from neuron # 116 to # 128. Statistics: p < 0.05, Student *t*-test with Bonferroni correction. Dotted lines represented lag times of -10, 0, and 10 msec (from left to right) after spiking of leading neuron # 116.

(B) As (A) for FC of another example congruent active pair.

(C) As (A) for FC of an example congruent inactive pair.

(D) As (A) for FC of an example incongruent active pair.

(E) As (A) for FC of an example incongruent inactive pair.

(F) As (A) for FC of an example non-memory pair.

(G) FR of various types of neuronal pairs in hit trials. Statistics: Tw-ANOVA-md, for the comparison between types of neuronal pairs, F (4,15) = 66.80, p = 2.24 × 10^−9^. Error bars represent mean ± SEM.

(H) FC density in CR trials controlling for FR, as Figure 3F. Statistics: Two-way ANOVA, for the comparison between pair types: F (3,3) = 3.43, p = 0.035; between FR levels: F (1,3) = 102.87, p = 9.32 × 10^−10^.

(I) FC density in neuronal pairs with different combinations of FR modulations during delay relative to baseline period. D represents decreased activity and I represents increased activity. Statistics: Two-way ANOVA, for the comparison between types of neuronal pairs, F (4,8) = 5.28, p = 0.0015; between combinations of FR modulations, F (2,8) = 30.61, p = 6.30 × 10^−9^.

(J) The proportion of following mPFC neurons capable of encoding STM information, with memory (black) or non-memory (white) mPFC neurons as the leading neuron, as Figure 3I. Statistics: p = 1.16 × 10^−16^, chi-square test.

**Figure S6. Projection-specific optogenetic manipulation. Related to Figure 4.**

(A) Performance in *d’* of mice in performing DRT, for optogenetic suppression of delay-period activity of mPFC-aAIC projections (green) and control (black) groups. Statistics: F (1,21) = 11.69, p = 0.0026, Tw-ANOVA-md for learning-day performance, n = 12 and 11 mice for NpHR and eYFP groups, respectively; p = 0.46 for laser off/on in the well-trained phase, Mann-Whitney *U* test, n = 8 and 7 mice for NpHR and eYFP groups, respectively. Error bars: mean ± SEM.

(B) Lick efficiency, following suppression of mPFC-aAIC projections. Statistics: F (1,21) = 6.36, p = 0.020, Tw-ANOVA-md for learning-day performance; p = 0.28 for laser off/on in the well-trained phase, Mann-Whitney *U* test. Number of mice as in (A).

(C) Proportion of aborted trials, following suppression of mPFC-aAIC projections. Statistics: F (1,21) = 0.037, p = 0.85, Tw-ANOVA-md for learning-day performance; p = 0.26 for laser off/on in the well-trained phase, Mann-Whitney *U* test. Number of mice as in (A).

(D) Accumulated histology images showing spatial extension of virus CaMKII_α_-NpHR and locations of optical fibers in mice in Figures 4C and 4E.

(E) Performance in *d’* of mice in performing DRT, for optogenetic activation of delay-period activity of mPFC-aAIC projections (blue) and control (black) groups. Statistics: F (1,12) = 10.48, p = 0.0071, Tw-ANOVA-md for learning-day performance, n = 7 mice for ChR2 and mCherry groups; p = 0.45 for laser off/on in the well-trained phase, Mann-Whitney *U* test, n = 7 and 6 mice for ChR2 and mCherry groups, respectively.

(F) As (B) for activation of mPFC-aAIC projections. Statistics: F (1,12) = 5.63, p = 0.035, Tw-ANOVA-md for learning-day performance; p = 0.79 for laser off/on in the well-trained phase, Mann-Whitney *U* test. Number of mice as in (E).

(G) As (C) for activating mPFC-aAIC projections. Statistics: F (1,12) = 6.16, p = 0.029, Tw-ANOVA-md for learning-day performance; p = 0.84 for laser off/on in the well-trained phase, Mann-Whitney *U* test. Number of mice as in (E).

(H) Accumulated histology images showing spatial extension of virus CaMKII_α_-ChR2 and locations of optical fibers in mice in Figures 4H and 4J.

**Figure S7. Simultaneous projection-specific optogenetic manipulation and recording in aAIC. Related to Figures 5 and 6.**

(A) Raster and FR of example aAIC neurons with suppressed (top) and activated (bottom) activity by projection-specific optogenetic suppression. Top green bars indicated for the laser-delivery period.

(B) The proportion (top) and normalized FR (bottom) of aAIC neurons with different modulations.

(C) Memory-coding ability of transient (left, F (1,337) = 5.81, p = 0.016) and sustained (right, F (1,64) = 2.14, p = 0.15) aAIC neurons, for optogenetic suppression of delay-period activity of mPFC-aAIC projections (green) and control (black) groups. Statistics: Tw-ANOVA-md. For transient neurons: n = 208 and 131 neurons for control and suppression groups, respectively; for sustained neurons: n = 50 and 16 neurons for control and suppression groups, respectively. Error bars represent mean ± SEM.

(D) Proportion of transient (left, p = 0.0070) and sustained (right, p = 6.78 × 10^−4^) aAIC neurons, following suppression of mPFC-aAIC projections. Statistics: chi-square test.

(E) Memory-coding ability of NoGo-preferred aAIC neurons, following suppression of mPFC-aAIC projections. Statistics: Tw-ANOVA-md, F (1,188) = 0.49, p = 0.48. n = 114 and 76 neurons for control and suppression groups, respectively. Error bars represent mean ± SEM.

(F) Proportion of memory aAIC neurons during the delay period (bin size: 1 sec), following suppression of mPFC-aAIC projections. Statistics: p = 0.0021, 3.07 × 10^−4^, 1.81 × 10^−4^, and 7.14 × 10^−5^ over the time course, chi-square test with Bonferroni correction. n = 437 and 345 neurons for control and suppression groups, respectively.

(G) Raster and FR of example aAIC neurons with optogenetic excitation of prefrontal projections. Top blue bars indicated the laser-delivery period.

(H) The proportion (top) and normalized FR (bottom) of activated aAIC neurons.

(I) Memory-coding ability of Go-preferred (left, F (1,311) = 0.36, p = 0.55) and NoGo-preferred (right, F (1,198) = 0.55, p = 0.46) aAIC neurons, for optogenetic excitation of delay-period activity of mPFC-aAIC projections (blue) and control (black) groups. Statistics: Tw-ANOVA-md. For Go-preferred neurons: n = 144 and 169 neurons for control and excitation groups, respectively; for NoGo-preferred neurons: n = 114 and 86 neurons for control and excitation groups, respectively. Error bars represent mean ± SEM.

(J) Proportion of transient (left, p = 0.014) and sustained (right, p = 0.99) aAIC neurons, following excitation of mPFC-aAIC projections. Statistics: chi-square test.

(K) Memory-coding ability of sustained aAIC neurons, following excitation of mPFC-aAIC projections. Statistics: Tw-ANOVA-md, F (1,91) = 4.10, p = 0.046. n = 50 and 43 neurons for control and excitation groups, respectively.

(L) Proportion of memory aAIC neurons during delay (bin size: 1 sec), following excitation of mPFC-aAIC projections. Statistics: p = 2.14, 0.24, 0.78, and 0.011 over the time course, chi-square test with Bonferroni correction. n = 437 and 377 neurons for control and excitation groups, respectively.

**Figure S8. STM-encoding ability of FC and efficacy of information transfer of aAIC memory neurons. Related to Figure 7.**

(A) STM-encoding ability of FC in congruent inactive (left, F (2,3) = 5.41, p = 4.75 × 10^−3^) and incongruent inactive (right, F (2,3) = 13.86, p = 2.03 × 10^−6^) pairs of aAIC neurons, following optogenetic suppression (green) or activation (blue) of mPFC-aAIC projections. Statistics: Two-way ANOVA. Error bars represent mean ± SEM.

(B) Difference in FR between different sample odors for FC pairs in congruent active (left, F (2,3) = 0.17, p = 0.84), congruent inactive (middle, F (2,3) = 0.33, p = 0.72), and incongruent inactive (right, F (2,3) = 0.59, p = 0.55) groups, following projection-specific manipulation. Statistics: Two-way ANOVA.

(C) FR of leading neurons in the trial type with significant FC in mPFC. Statistics: ***p < 0.001, bootstrap test. Error bars indicate 95% CI from bootstrap of 1000 times.

(D) Change in STM-encoding ability of following mPFC neurons after removing FCSPs from leading mPFC neurons, controlling for FR of leading neurons. At 8-10 Hz (left), for the comparison between types of leading neurons, F (1,3) = 29.43, p = 1.17 × 10^−7^; between number of leading neurons, F (3,3) = 3.96, p = 0.0086. At 10-12 Hz (right), between types of leading neurons, F (1,3) = 51.98, p = 3.57 × 10^−12^; between number of leading neurons, F (3,3) = 3.75, p = 0.011. Statistics: Two-way ANOVA.

(E) As (C) for FC pairs across regions (left, p = 0.37 and 0.27 for 1 and 2 leading neurons with removed FCSP spikes, respectively) and in aAIC (right, p = 0.18 and 0.44 for 1 and 2 leading neurons with removed FCSP spikes, respectively). Statistics: bootstrap test.

(F) Left, change in STM-encoding ability of following aAIC neurons with the removal of FCSPs from aAIC neurons, following projection-specific manipulation. For memory leading neurons: F (2,502) = 3.57, p = 0.0289; for non-memory leading neurons: F (2,338) = 0.23, p = 0.80. Right, activity of leading neurons in the trial type with significant FC, following projection-specific manipulation. For memory leading neurons: F (2,554) = 15.18, p = 3.82 × 10^−7^; for non-memory leading neurons: F (2,374) = 1.96, p = 0.14. Statistics: One-way ANOVA.

