## Supplemental Figures for "Prefrontal projections modulate recurrent circuitry in insular cortex to support short-term memory"

**Figure S1**

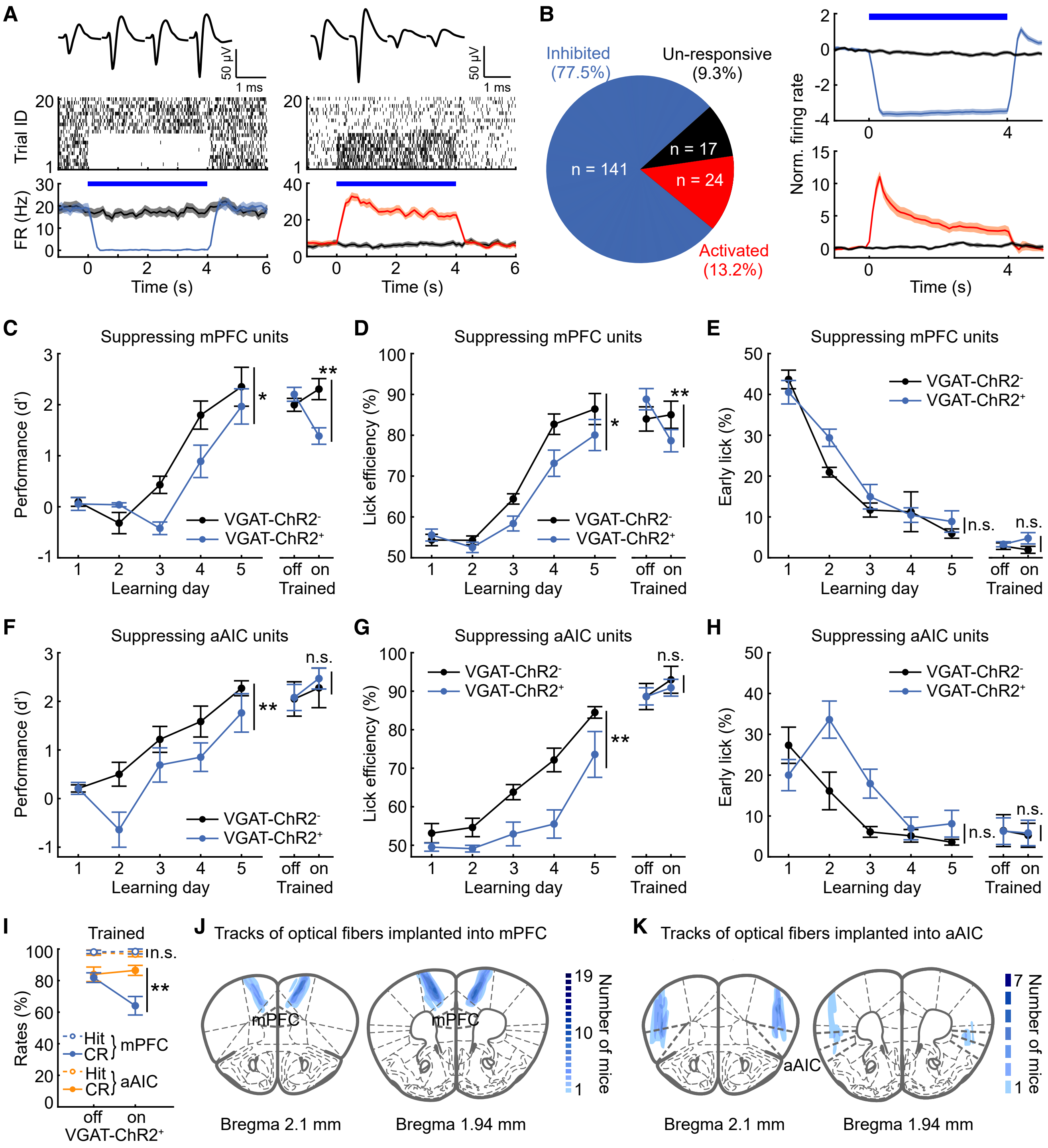
A
50 μV
1 ms
Trial ID
FR (Hz)
Time (s)
B
Inhibited (77.5%)
Un-responsive (9.3%)
Activated (13.2%)
n = 141
n = 17
n = 24
Norm. firing rate
Time (s)
C
Suspressing mPFC units
Performance (d')
Learning day
Trained
VGAT-ChR2-
VGAT-ChR2+
D
Suppressing mPFC units
Lick efficiency (%)
E
Suppressing mPFC units
Early lick (%)
n.s.
F
Suppressing aAIC units
G
Suppressing aAIC units
H
Suppressing aAIC units
I
Trained
Rates (%)
Hit
CR
mPFC
aAIC
VGAT-ChR2+
J
Tracks of optical fibers implanted into mPFC
Bregma 2.1 mm
Bregma 1.94 mm
Number of mice
K
Tracks of optical fibers implanted into aAIC

**Figure S1. Information associated with optogenetic suppression. Related to Figure 1.**

(A) Waveforms (first row), raster plots (second row), and PSTH plots (third row) of two example mPFC neurons with different modulations by laser. (B) The proportion (left) and population averaged normalized FR (right) of mPFC neurons with different modulations. (C) Performance in d' of mice in performing DRT, for optogenetic suppression of mPFC delay-period activity (blue) and control (black) groups. Statistics: $F(1,19) = 6.07$, $p = 0.023$, Tw-ANOVA-md for learning-day performance, $n = 11$ and 10 mice for VGAT-ChR2$^{+}$ and VGAT-ChR2$^{-}$ groups, respectively; $p = 0.0019$ for laser off/on in the well-trained phase, Mann-Whitney U test, $n = 8$ mice for both groups. Error bars indicate mean ± SEM. (D) Lick efficiency, following suppression of mPFC. Statistics: $F(1,19) = 4.75$, $p = 0.042$, Tw-ANOVA-md for learning-day performance; $p = 0.0030$ for laser off/on in the well-trained phase, Mann-Whitney U test. Number of mice as in (C). (E) Proportion of aborted (early lick) trials, following suppression of mPFC. Statistics: $F(1,19) = 1.13$, $p = 0.30$, Tw-ANOVA-md for learning-day performance; $p = 0.12$ for laser off/on in the well-trained phase, Mann-Whitney U test. Number of mice as in (C). (F) As (C) for suppressing aAIC delay-period activity. Statistics: $F(1,13) = 15.77$, $p = 0.0016$, Tw-ANOVA-md for learning-day performance, $n = 7$ and 8 mice for VGAT-ChR2$^{+}$ and VGAT-ChR2$^{-}$ groups, respectively; $p = 0.86$ for laser off/on in the well-trained phase, Mann-Whitney U test, $n = 8$ mice for both groups. (G) As (D) for suppression of aAIC. Statistics: $F(1,13) = 14.59$, $p = 0.0021$, Tw-ANOVA-md for learning-day performance; $p = 0.57$ for laser off/on in the well-trained phase, Mann-Whitney U test. Number of mice as in (F). (H) As (E) for suppressing aAIC. Statistics: $F(1,13) = 4.28$, $p = 0.059$, Tw-ANOVA-md for learning-day performance; $p = 0.46$ for laser off/on in the well-trained phase, Mann-Whitney U test. Number of mice as in (F). (I) Hit and CR rates of mice in performing DRT in the well-trained phase, following suppression of mPFC (blue) or aAIC (orange) delay-period activity. Statistics: Mann-Whitney U test, $p = 0.49$ and 0.0016 for the hit and CR rate, respectively. $n = 8$ mice for suppressing mPFC and aAIC groups, respectively. (J) Accumulated histology images showing locations of optical fibers on top of mPFC in mice in Figures 1B and 1E. (K) Accumulated histology images showing locations of optical fibers on top of aAIC in mice in Figures 1G and 1J.

#### Figure S2

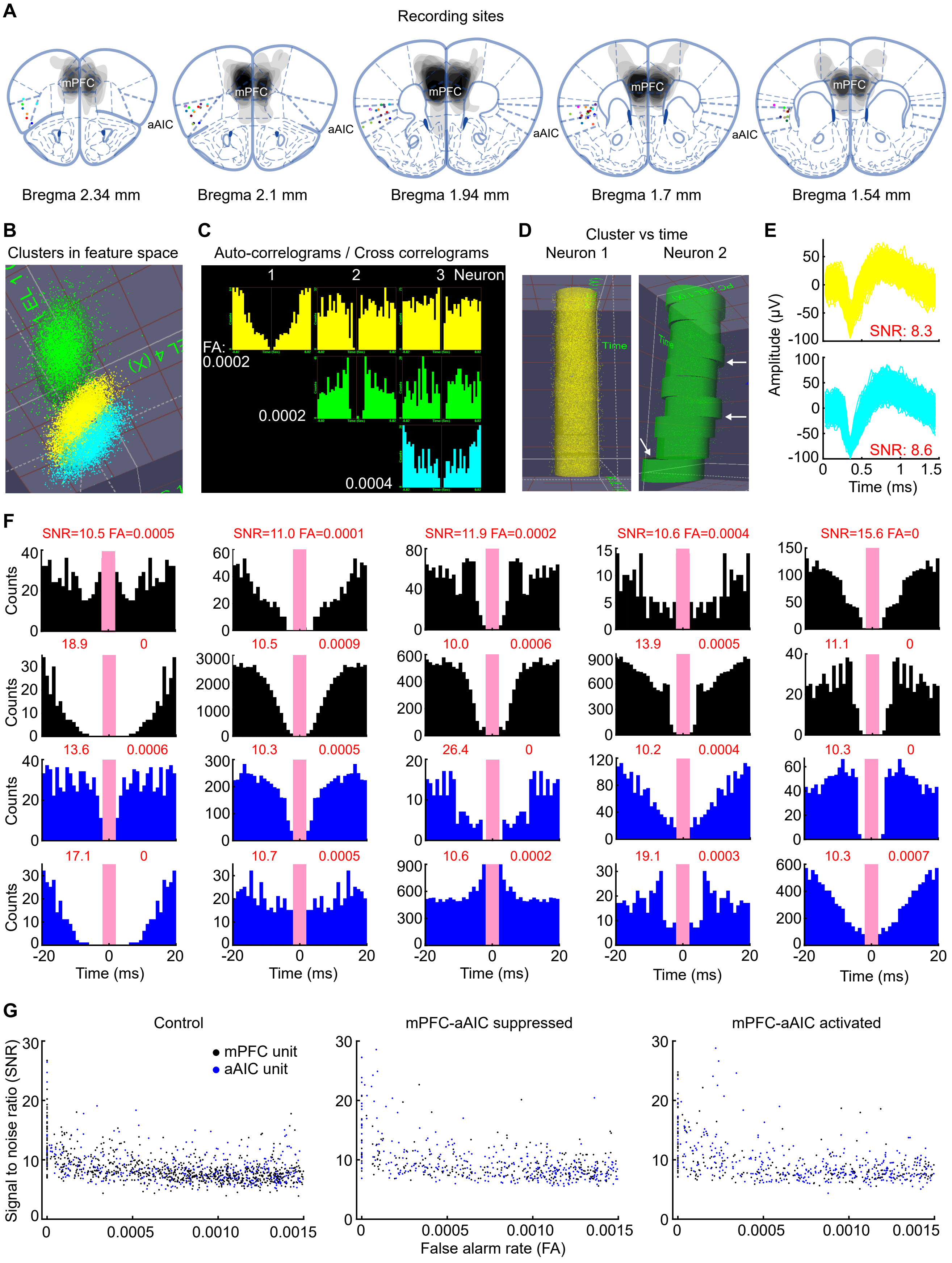

A
Recording sites
mPFC
aAIC
Bregma 2.34 mm
Bregma 2.1 mm
Bregma 1.94 mm
Bregma 1.7 mm
Bregma 1.54 mm
B
Clusters in feature space
C
Auto-correlograms / Cross correlograms
1
2
3 Neuron
FA:
0.0002
0.0002
0.0004
D
Cluster vs time
Neuron 1
Neuron 2
Time
E
SNR: 8.3
SNR: 8.6
Amplitude (μV)
Time (ms)
F
SNR=10.5 FA=0.0005
SNR=11.0 FA=0.0001
SNR=11.9 FA=0.0002
SNR=10.6 FA=0.0004
SNR=15.6 FA=0
18.9 0
10.5 0.0009
10.0 0.0006
13.9 0.0005
11.1 0
13.6 0.0006
10.3 0.0005
26.4 0
10.2 0.0004
10.3 0
17.1 0
10.7 0.0005
10.6 0.0002
19.1 0.0003
10.3 0.0007
Counts
Time (ms)
G
Control
mPFC-aAIC suppressed
mPFC-aAIC activated
mPFC unit
aAIC unit
Signal to noise ratio (SNR)
False alarm rate (FA)

**Figure S2. Spike sorting. Related to Figure 2.**

(A) Accumulated images showing unilateral extracellular recording sites in aAIC based on electricity lesion after recording for Figures 2, 5, and 6, with the dots of different color indicated for recording site from different mice. Shaded area indicated accumulated virus expression in mPFC in Figures 5 and 6. (B) Three example clusters of spikes in the feature space based on PCA of all spikes from one tetrode. (C) Spike autocorrelogram and cross-correlogram for clusters shown in (B).
(D) Distribution of the same clusters in (B) cross time, showing the stability of the recorded neuron. White arrows in Neuron 2 indicated unstable examples. This type of unstable neurons were excluded from further analysis. (E) Waveforms of two qualified neurons included for further analysis in (B). Signal to noise ratio (SNR): mean peak amplitude of all spikes divided by standard deviation during baseline. (F) Autocorrelograms for example neurons. False alarm (FA) rate is the ratio of spikes within inter-spike-interval less than 2 msec. Black for mPFC and blue for aAIC neurons.
(G) Scatter plots of SNR and FA rate of all recorded neurons.

#### Figure S3

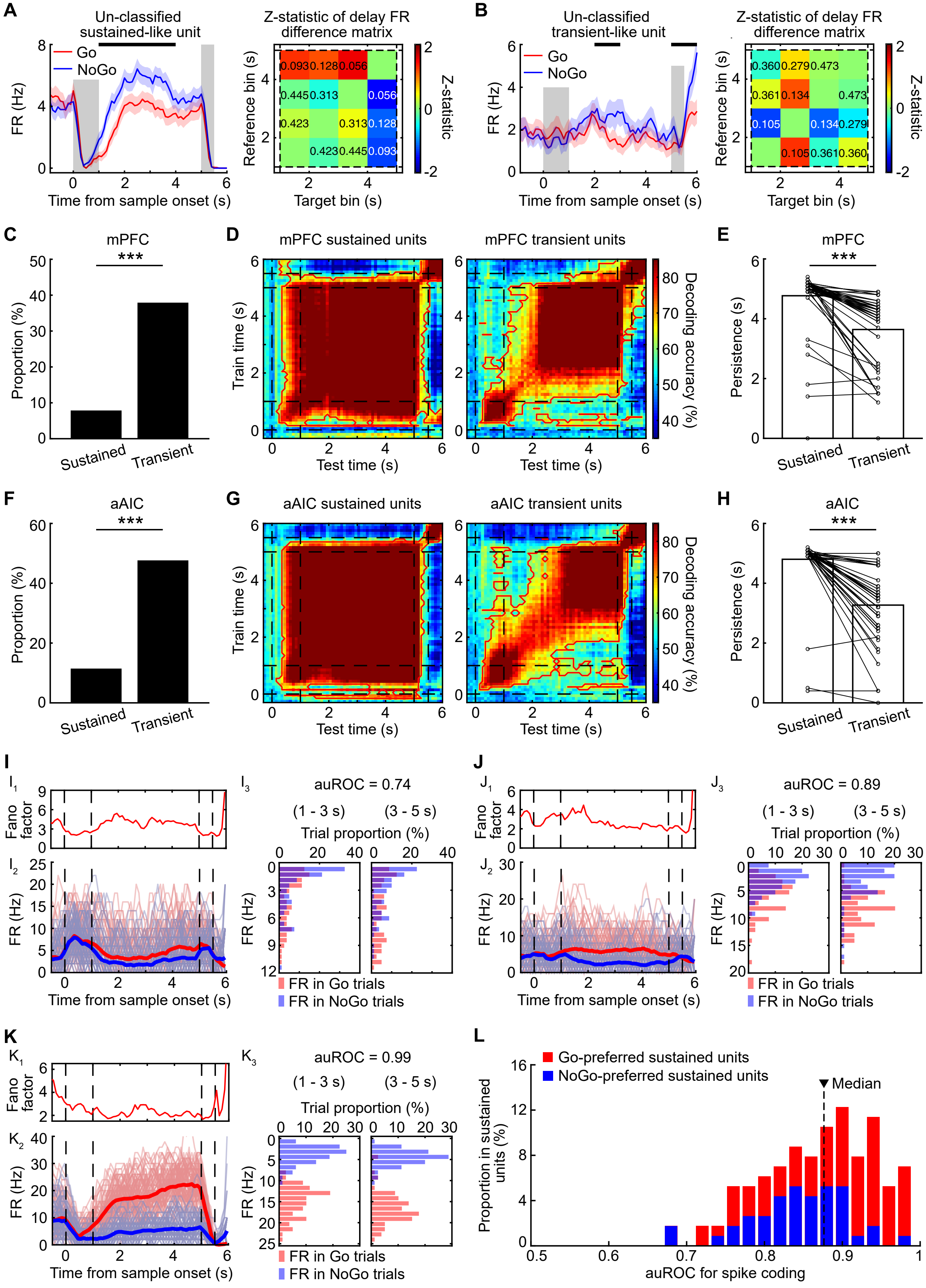

A
Un-classified sustained-like unit
Go
NoGo
FR (Hz)
Time from sample onset (s)
Z-statistic of delay FR difference matrix
Reference bin (s)
Target bin (s)
Z-statistic
B
Un-classified transient-like unit
C
mPFC
Proportion (%)
Sustained
Transient
D
mPFC sustained units
mPFC transient units
Train time (s)
Test time (s)
Decoding accuracy (%)
E
mPFC
Persistence (s)
F
aAIC
G
aAIC sustained units
aAIC transient units
H
aAIC
I
auROC = 0.74
(1 - 3 s)
(3 - 5 s)
Trial proportion (%)
Fano factor
FR in Go trials
FR in NoGo trials
J
auROC = 0.89
K
auROC = 0.99
L
Go-preferred sustained units
NoGo-preferred sustained units
Median
Proportion in sustained units (%)
auROC for spike coding

**Figure S3. Transient and sustained neurons in mPFC and aAIC in DRT. Related to Figure 2.**

(A) An example unclassified neuron exhibiting 'sustained'-like selectivity. Left, PSTH plot. Statistics: $p < 0.05$, Mann-Whitney U test with Bonferroni correction. Right, Z-statistic of FR difference matrix during delay. Statistics: permutation test, with p values indicated by numbers. Because of the large P value between the activity of different time bins, these neurons (n = 7/4 neurons for mPFC/aAIC) were not classified into sustained or transient group. (B) An example un-classified 'transient'-like neuron. Note these neurons (n = 30/14 for mPFC/aAIC) were not classified into transient group. (C) Proportion of sustained (7.84%) and transient (37.87%) mPFC neurons. Statistics: $p = 0$, chi-square test. (D) CTD results with sustained (left) and transient (right) mPFC neurons, as Figure 2G. (E) Persistence in memory-coding ability of transient and sustained mPFC neurons, indicating the stability of information maintenance. Dots indicated for coding persistence of different training time point, which was calculated as the duration of significant decoding from the onset of sample odor to the offset of test odor. Statistics: $p = 2.89 \times 10^{-10}$, Wilcoxon signed-rank test. (F) Proportion of sustained (11.44%) and transient (47.60%) aAIC neurons. Statistics: $p = 0$, chi-square test. (G) Same as (D) for sustained (left) and transient (right) aAIC neurons. (H) Same as (E) for transient and sustained aAIC neurons. Statistics: $p = 4.87 \times 10^{-10}$, Wilcoxon signed-rank test. (I) Cross-trial FR of an example sustained neuron, with auROC value 0.74. (I1) Fano Factor in preferred Go trials. (I2) FR of each single trials. (I3) Distribution of FR of all trials. (J) As (I) for another sustained neuron, with auROC value 0.89. (K) As (I) for another sustained neuron, with auROC value 0.99. (L) Distribution of memory-coding ability of all sustained neurons (n = 114 neurons).

**Figure S4**

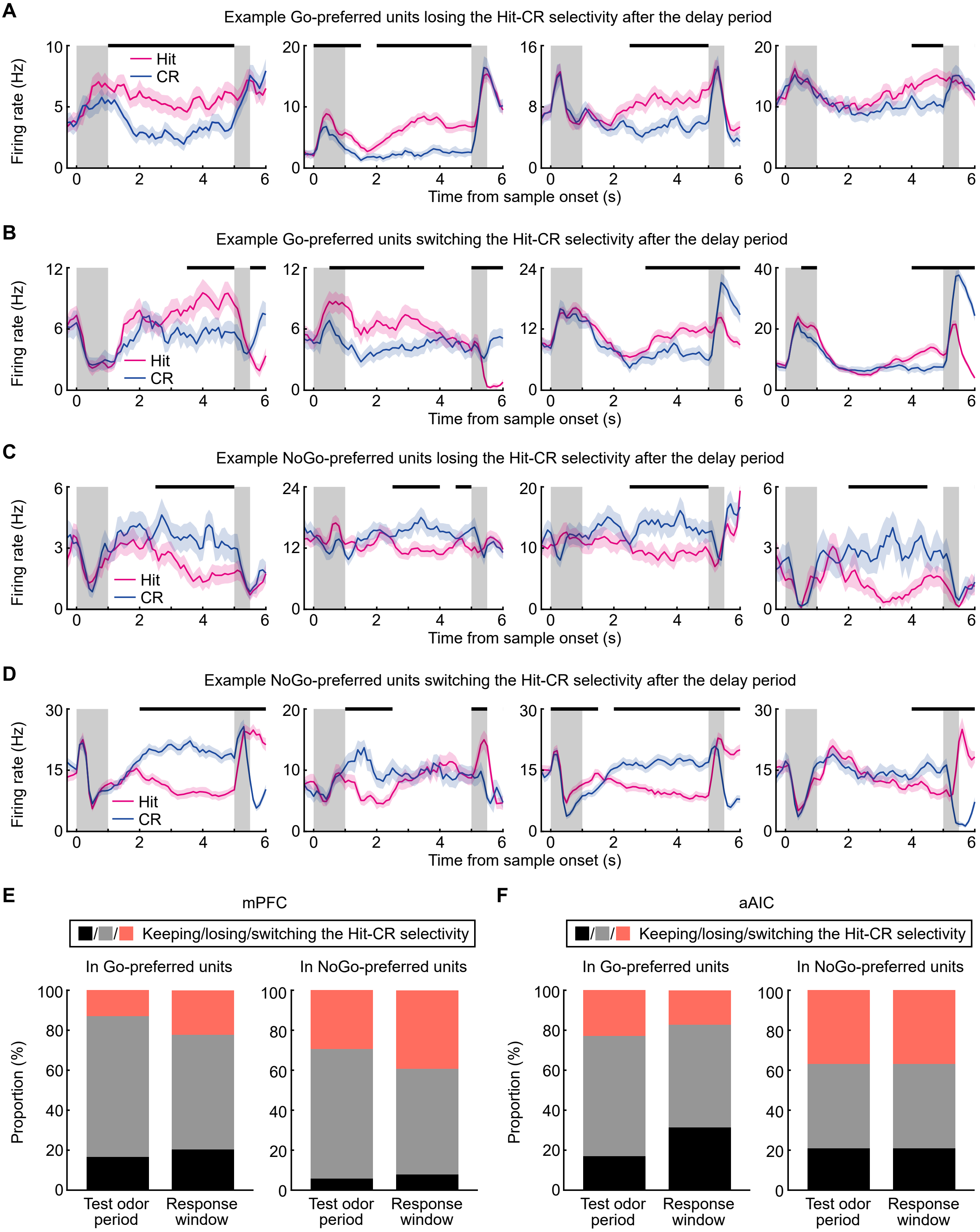


**Figure S4. Selectivity of memory neurons during delay and action pariods. Related to Figure 2.**

(A) FR of example Go-preferred neurons, losing selectivity for hit and CR trials after delay. (B) FR of example Go-preferred neurons, switching selectivity for hit and CR trials after delay. (C) As (A) for NoGo-preferred neurons. (D) As (B) for NoGo-preferred neurons. (E) Proportion of neurons keeping, losing, and switching selectivity in the Go-preferred (left) and NoGo-preferred (right) mPFC neurons. (F) As (E) for aAIC.

#### Figure S5

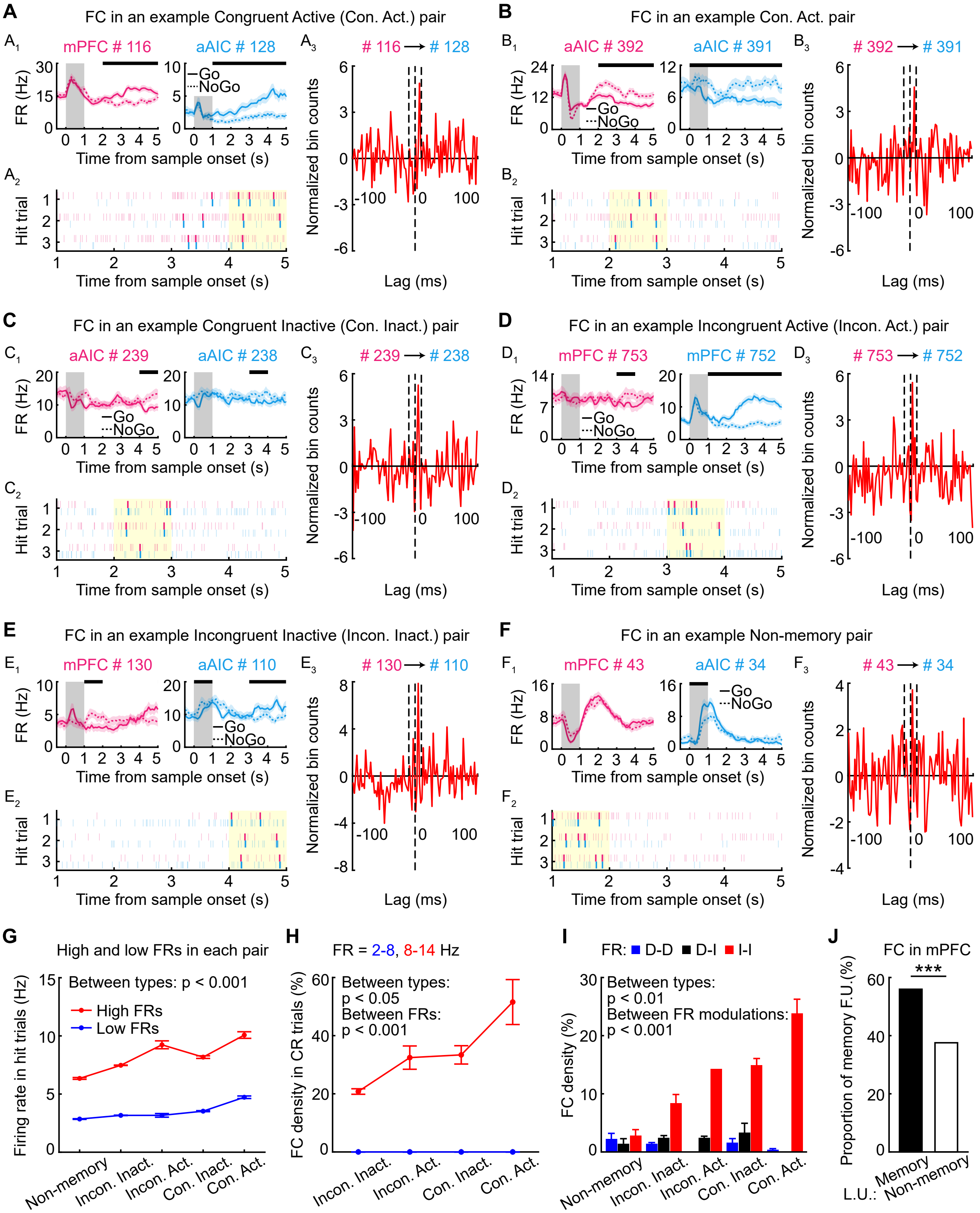
A
FC in an example Congruent Active (Con. Act.) pair
mPFC # 116
aAIC # 128
# 116 → # 128
B
FC in an example Con. Act. pair
aAIC # 392
aAIC # 391
# 392 → # 391
C
FC in an example Congruent Inactive (Con. Inact.) pair
aAIC # 239
aAIC # 238
# 239 → # 238
D
FC in an example Incongruent Active (Incon. Act.) pair
mPFC # 753
mPFC # 752
# 753 → # 752
E
FC in an example Incongruent Inactive (Incon. Inact.) pair
mPFC # 130
aAIC # 110
# 130 → # 110
F
FC in an example Non-memory pair
mPFC # 43
aAIC # 34
# 43 → # 34
FR (Hz)
Time from sample onset (s)
Hit trial
Normalized bin counts
Lag (ms)
Go
NoGo
G
High and low FRs in each pair
Between types: p < 0.001
High FRs
Low FRs
Firing rate in hit trials (Hz)
Non-memory
Incon. Inact.
Incon. Act.
Con. Inact.
Con. Act.
H
FR = 2-8, 8-14 Hz
Between types: p < 0.05
Between FRs: p < 0.001
FC density in CR trials (%)
I
FR: D-D D-I I-I
Between types: p < 0.01
Between FR modulations: p < 0.001
FC density (%)
J
FC in mPFC
***
Proportion of memory F.U.(%)
L.U.: Memory
Non-memory

**Figure S5. Preferential functional coupling between memory neurons. Related to Figure 3.**

(A) FC of an example congruent active pair (mPFC neuron # 116 and aAIC # 128). (A1) PSTH plots of the two example neurons. (A2) Raster plot showing spike timestamps in three example hit trials. Pink and blue bars indicated for spikes of mPFC neuron # 116 and aAIC # 128, respectively. Thick line pairs indicated FCSPs. Yellow region represents the period in which spike trains were used in the cross-correlogram analysis. (A3) Z-scored SSCC between spike trains of the two neurons. A prominent displaced peak on the right side indicated the directional FC from neuron # 116 to # 128. Statistics: $p < 0.05$, Student t-test with Bonferroni correction. Dotted lines represented lag times of -10, 0, and 10 msec (from left to right) after spiking of leading neuron # 116. (B) As (A) for FC of another example congruent active pair. (C) As (A) for FC of an example congruent inactive pair. (D) As (A) for FC of an example incongruent active pair. (E) As (A) for FC of an example incongruent inactive pair. (F) As (A) for FC of an example non-memory pair. (G) FR of various types of neuronal pairs in hit trials. Statistics: Tw-ANOVA-md, for the comparison between types of neuronal pairs, $F (4,15) = 66.80$, $p = 2.24 \times 10^{-9}$. Error bars represent mean ± SEM. (H) FC density in CR trials controlling for FR, as Figure 3F. Statistics: Two-way ANOVA, for the comparison between pair types: $F (3,3) = 3.43$, $p = 0.035$; between FR levels: $F (1,3) = 102.87$, $p = 9.32 \times 10^{-10}$. (I) FC density in neuronal pairs with different combinations of FR modulations during delay relative to baseline period. D represents decreased activity and I represents increased activity. Statistics: Two-way ANOVA, for the comparison between types of neuronal pairs, $F (4,8) = 5.28$, $p = 0.0015$; between combinations of FR modulations, $F (2,8) = 30.61$, $p = 6.30 \times 10^{-9}$. (J) The proportion of following mPFC neurons capable of encoding STM information, with memory (black) or non-memory (white) mPFC neurons as the leading neuron, as Figure 3I. Statistics: $p = 1.16 \times 10^{-16}$, chi-square test.

**Figure S6**

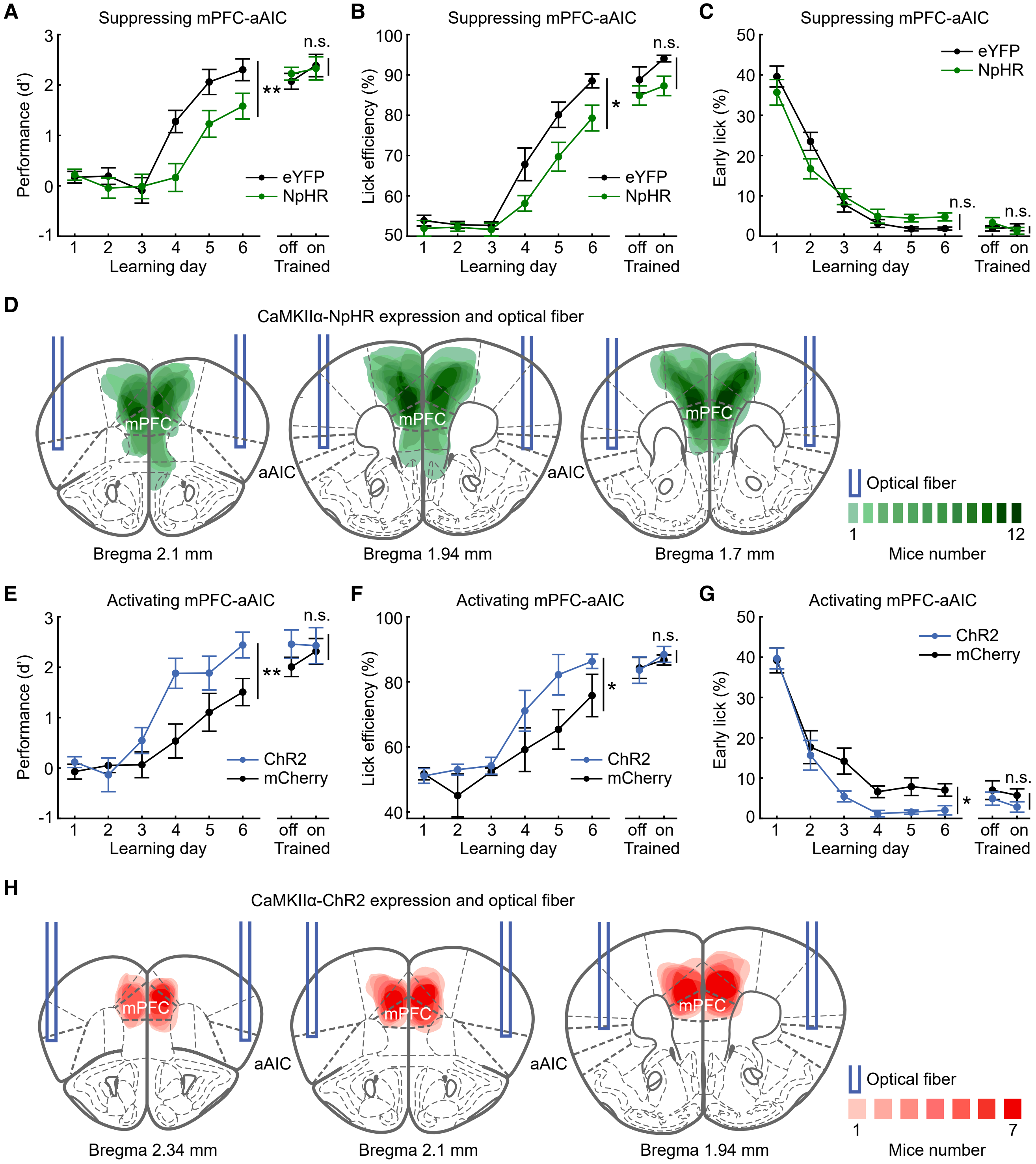
A
Suppressing mPFC-aAIC
Performance (d')
Learning day
Trained
off on
eYFP
NpHR
**
n.s.
B
Suppressing mPFC-aAIC
Lick efficiency (%)
*
n.s.
C
Suppressing mPFC-aAIC
Early lick (%)
n.s.
D
CaMKIIα-NpHR expression and optical fiber
mPFC
aAIC
Bregma 2.1 mm
Bregma 1.94 mm
Bregma 1.7 mm
Optical fiber
1
12
Mice number
E
Activating mPFC-aAIC
ChR2
mCherry
F
Activating mPFC-aAIC
G
Activating mPFC-aAIC
H
CaMKIIα-ChR2 expression and optical fiber
Bregma 2.34 mm
Bregma 2.1 mm
Bregma 1.94 mm
7

**Figure S6. Projection-specific optogenetic manipulation. Related to Figure 4.**

(A) Performance in d’ of mice in performing DRT, for optogenetic suppression of delay-period activity of mPFC-aAIC projections (green) and control (black) groups. Statistics: $F (1,21) = 11.69$, $p = 0.0026$, Tw-ANOVA-md for learning-day performance, n = 12 and 11 mice for NpHR and eYFP groups, respectively; $p = 0.46$ for laser off/on in the well-trained phase, Mann-Whitney U test, n = 8 and 7 mice for NpHR and eYFP groups, respectively. Error bars: mean ± SEM. (B) Lick efficiency, following suppression of mPFC-aAIC projections. Statistics: $F (1,21) = 6.36$, $p = 0.020$, Tw-ANOVA-md for learning-day performance; $p = 0.28$ for laser off/on in the well-trained phase, Mann-Whitney U test. Number of mice as in (A). (C) Proportion of aborted trials, following suppression of mPFC-aAIC projections. Statistics: $F (1,21) = 0.037$, $p = 0.85$, Tw-ANOVA-md for learning-day performance; $p = 0.26$ for laser off/on in the well-trained phase, Mann-Whitney U test. Number of mice as in (A). (D) Accumulated histology images showing spatial extension of virus CaMKIIα-NpHR and locations of optical fibers in mice in Figures 4C and 4E. (E) Performance in d’ of mice in performing DRT, for optogenetic activation of delay-period activity of mPFC-aAIC projections (blue) and control (black) groups. Statistics: $F (1,12) = 10.48$, $p = 0.0071$, Tw-ANOVA-md for learning-day performance, n = 7 mice for ChR2 and mCherry groups; $p = 0.45$ for laser off/on in the well-trained phase, Mann-Whitney U test, n = 7 and 6 mice for ChR2 and mCherry groups, respectively. (F) As (B) for activation of mPFC-aAIC projections. Statistics: $F (1,12) = 5.63$, $p = 0.035$, Tw-ANOVA-md for learning-day performance; $p = 0.79$ for laser off/on in the well-trained phase, Mann-Whitney U test. Number of mice as in (E). (G) As (C) for activating mPFC-aAIC projections. Statistics: $F (1,12) = 6.16$, $p = 0.029$, Tw-ANOVA-md for learning-day performance; $p = 0.84$ for laser off/on in the well-trained phase, Mann-Whitney U test. Number of mice as in (E).
(H) Accumulated histology images showing spatial extension of virus CaMKIIα-ChR2 and locations of optical fibers in mice in Figures 4H and 4J.

### Figure S7

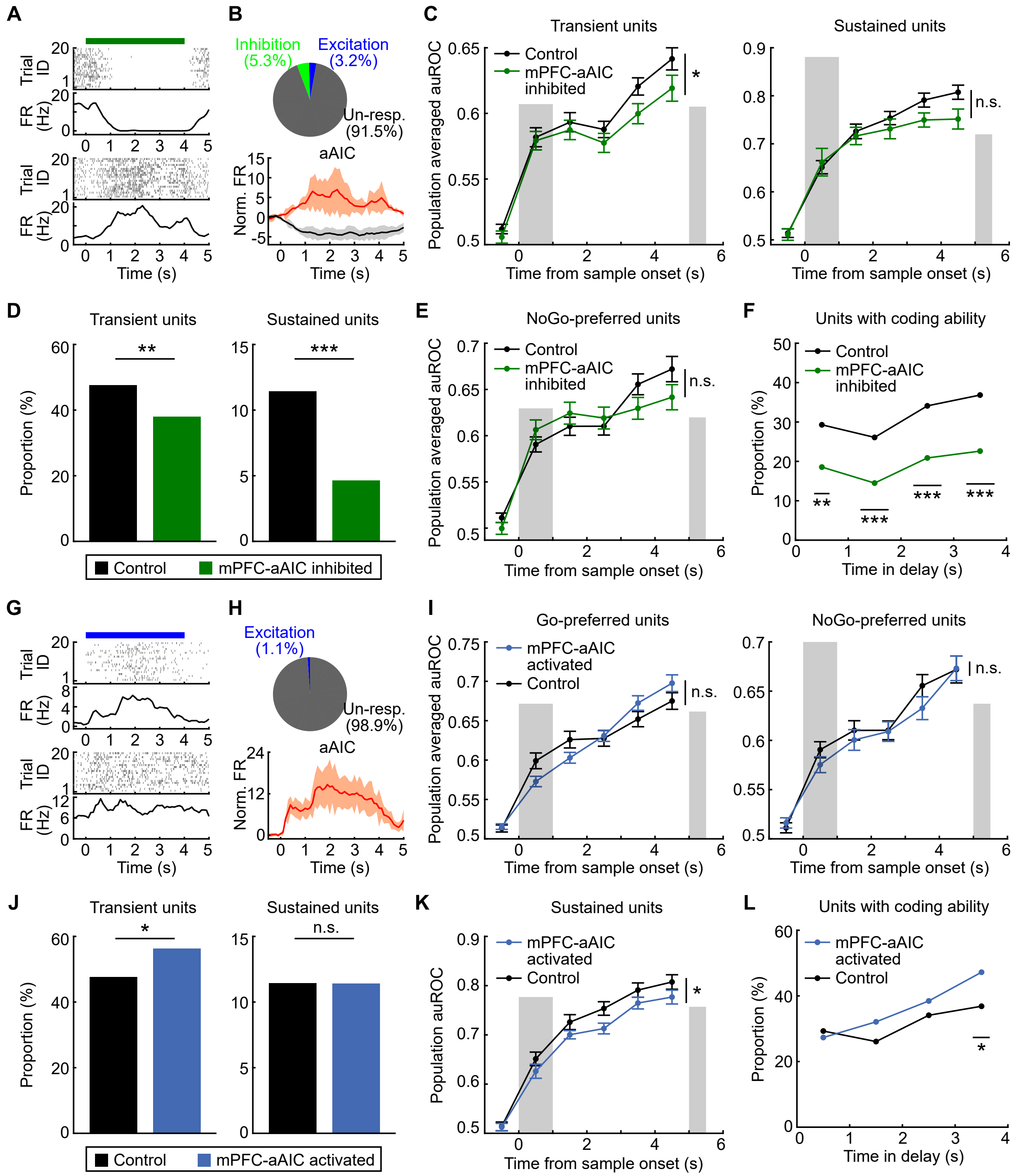

A
B
C
D
E
F
G
H
I
J
K
L
Inhibition (5.3%)
Excitation (3.2%)
Un-resp. (91.5%)
aAIC
Excitation (1.1%)
Un-resp. (98.9%)
Trial ID
FR (Hz)
Norm. FR
Time (s)
Transient units
Sustained units
NoGo-preferred units
Go-preferred units
Units with coding ability
Control
mPFC-aAIC inhibited
mPFC-aAIC activated
Population averaged auROC
Population auROC
Proportion (%)
Time from sample onset (s)
Time in delay (s)
n.s.
Control
mPFC-aAIC inhibited
Control
mPFC-aAIC activated

**Figure S7. Simultaneous projection-specific optogenetic manipulation and recording in aAIC. Related to Figures 5 and 6.**

(A) Raster and FR of example aAIC neurons with suppressed (top) and activated (bottom) activity by projection-specific optogenetic suppression. Top green bars indicated for the laser-delivery period. (B) The proportion (top) and normalized FR (bottom) of aAIC neurons with different modulations. (C) Memory-coding ability of transient (left, $F(1,337) = 5.81$, $p = 0.016$) and sustained (right, $F(1,64) = 2.14$, $p = 0.15$) aAIC neurons, for optogenetic suppression of delay-period activity of mPFC-aAIC projections (green) and control (black) groups. Statistics: Tw-ANOVA-md. For transient neurons: $n = 208$ and 131 neurons for control and suppression groups, respectively; for sustained neurons: $n = 50$ and 16 neurons for control and suppression groups, respectively. Error bars represent mean ± SEM. (D) Proportion of transient (left, $p = 0.0070$) and sustained (right, $p = 6.78 \times 10^{-4}$) aAIC neurons, following suppression of mPFC-aAIC projections. Statistics: chi-square test. (E) Memory-coding ability of NoGo-preferred aAIC neurons, following suppression of mPFC-aAIC projections. Statistics: Tw-ANOVA-md, $F(1,188) = 0.49$, $p = 0.48$. $n = 114$ and 76 neurons for control and suppression groups, respectively. Error bars represent mean ± SEM. (F) Proportion of memory aAIC neurons during the delay period (bin size: 1 sec), following suppression of mPFC-aAIC projections. Statistics: $p = 0.0021$, $3.07 \times 10^{-4}$, $1.81 \times 10^{-4}$, and $7.14 \times 10^{-5}$ over the time course, chi-square test with Bonferroni correction. $n = 437$ and 345 neurons for control and suppression groups, respectively. (G) Raster and FR of example aAIC neurons with optogenetic excitation of prefrontal projections. Top blue bars indicated the laser-delivery period. (H) The proportion (top) and normalized FR (bottom) of activated aAIC neurons. (I) Memory-coding ability of Go-preferred (left, $F(1,311) = 0.36$, $p = 0.55$) and NoGo-preferred (right, $F(1,198) = 0.55$, $p = 0.46$) aAIC neurons, for optogenetic excitation of delay-period activity of mPFC-aAIC projections (blue) and control (black) groups. Statistics: Tw-ANOVA-md. For Go-preferred neurons: $n = 144$ and 169 neurons for control and excitation groups, respectively; for NoGo-preferred neurons: $n = 114$ and 86 neurons for control and excitation groups, respectively. Error bars represent mean ± SEM. (J) Proportion of transient (left, $p = 0.014$) and sustained (right, $p = 0.99$) aAIC neurons, following excitation of mPFC-aAIC projections. Statistics: chi-square test. (K) Memory-coding ability of sustained aAIC neurons, following excitation of mPFC-aAIC projections. Statistics: Tw-ANOVA-md, $F(1,91) = 4.10$, $p = 0.046$. $n = 50$ and 43 neurons for control and excitation groups, respectively. (L) Proportion of memory aAIC neurons during delay (bin size: 1 sec), following excitation of mPFC-aAIC projections. Statistics: $p = 2.14$, 0.24, 0.78, and 0.011 over the time course, chi-square test with Bonferroni correction. $n = 437$ and 377 neurons for control and excitation groups, respectively.

**Figure S8**

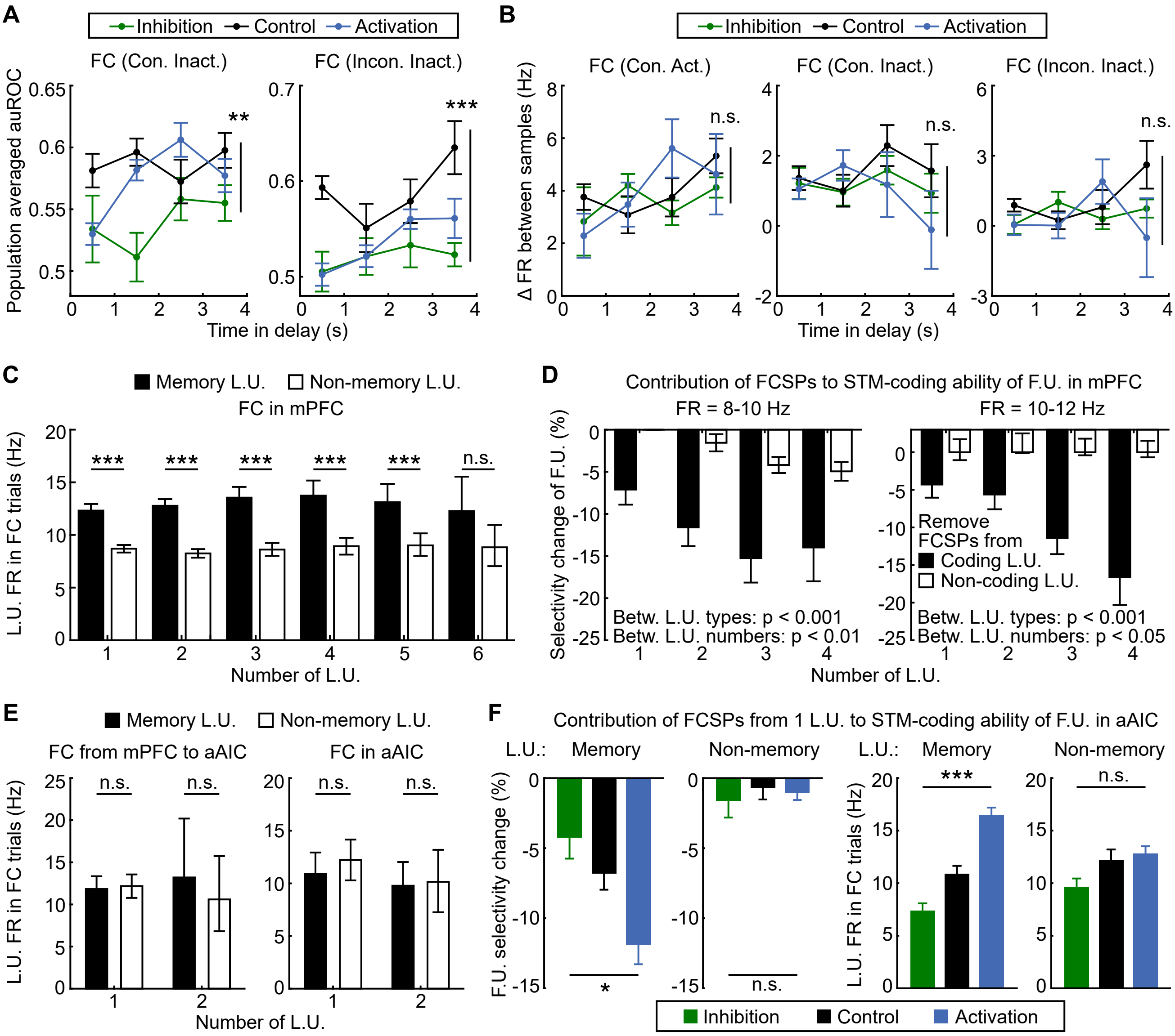
A
Inhibition
Control
Activation
FC (Con. Inact.)
FC (Incon. Inact.)
Population averaged auROC
Time in delay (s)
B
FC (Con. Act.)
FC (Con. Inact.)
FC (Incon. Inact.)
Δ FR between samples (Hz)
n.s.
C
Memory L.U.
Non-memory L.U.
FC in mPFC
L.U. FR in FC trials (Hz)
Number of L.U.
D
Contribution of FCSPs to STM-coding ability of F.U. in mPFC
FR = 8-10 Hz
FR = 10-12 Hz
Selectivity change of F.U. (%)
Remove FCSPs from
Coding L.U.
Non-coding L.U.
Betw. L.U. types: p < 0.001
Betw. L.U. numbers: p < 0.01
Betw. L.U. types: p < 0.001
Betw. L.U. numbers: p < 0.05
Number of L.U.
E
Memory L.U.
Non-memory L.U.
FC from mPFC to aAIC
FC in aAIC
L.U. FR in FC trials (Hz)
Number of L.U.
F
Contribution of FCSPs from 1 L.U. to STM-coding ability of F.U. in aAIC
L.U.: Memory
Non-memory
L.U.: Memory
Non-memory
F.U. selectivity change (%)
L.U. FR in FC trials (Hz)
Inhibition
Control
Activation

**Figure S8. STM-encoding ability of FC and efficacy of information transfer of aAIC memory neurons. Related to Figure 7.**

(A) STM-encoding ability of FC in congruent inactive (left, $F (2,3) = 5.41$, $p = 4.75 \times 10^{-3}$) and incongruent inactive (right, $F (2,3) = 13.86$, $p = 2.03 \times 10^{-6}$) pairs of aAIC neurons, following optogenetic suppression (green) or activation (blue) of mPFC-aAIC projections. Statistics: Two-way ANOVA. Error bars represent mean ± SEM. (B) Difference in FR between different sample odors for FC pairs in congruent active (left, $F (2,3) = 0.17$, $p = 0.84$), congruent inactive (middle, $F (2,3) = 0.33$, $p = 0.72$), and incongruent inactive (right, $F (2,3) = 0.59$, $p = 0.55$) groups, following projection-specific manipulation. Statistics: Two-way ANOVA. (C) FR of leading neurons in the trial type with significant FC in mPFC. Statistics: ***$p < 0.001$, bootstrap test. Error bars indicate 95% CI from bootstrap of 1000 times. (D) Change in STM-encoding ability of following mPFC neurons after removing FCSPs from leading mPFC neurons, controlling for FR of leading neurons. At 8-10 Hz (left), for the comparison between types of leading neurons, $F (1,3) = 29.43$, $p = 1.17 \times 10^{-7}$; between number of leading neurons, $F (3,3) = 3.96$, $p = 0.0086$. At 10-12 Hz (right), between types of leading neurons, $F (1,3) = 51.98$, $p = 3.57 \times 10^{-12}$; between number of leading neurons, $F (3,3) = 3.75$, $p = 0.011$. Statistics: Two-way ANOVA. (E) As (C) for FC pairs across regions (left, $p = 0.37$ and 0.27 for 1 and 2 leading neurons with removed FCSP spikes, respectively) and in aAIC (right, $p = 0.18$ and 0.44 for 1 and 2 leading neurons with removed FCSP spikes, respectively). Statistics: bootstrap test. (F) Left, change in STM-encoding ability of following aAIC neurons with the removal of FCSPs from aAIC neurons, following projection-specific manipulation. For memory leading neurons: $F (2,502) = 3.57$, $p = 0.0289$; for non-memory leading neurons: $F (2,338) = 0.23$, $p = 0.80$. Right, activity of leading neurons in the trial type with significant FC, following projection-specific manipulation. For memory leading neurons: $F (2,554) = 15.18$, $p = 3.82 \times 10^{-7}$; for non-memory leading neurons: $F (2,374) = 1.96$, $p = 0.14$. Statistics: One-way ANOVA.
